# Gain-of-function mutations in 1-phosphofructokinas*e (FruK)* re-establish glycolysis in bacteria

**DOI:** 10.64898/2026.11.13.766522

**Authors:** Beau Dronsella, Jan Zarzycki, Daniel Bonder, Irina Piven, Stefan Hristov, Ehsan Salehabadi, Lucía Torregrosa-Barragán, Ari Satanowski, Luca Schulz, Arren Bar-Even, Tobias. J. Erb, Steffen N. Lindner

## Abstract

Secondary (promiscuous) activities provide enzymes with a reservoir of latent catalytic functions, which can be leveraged during selective pressure. Here we retrace the re-emergence of glycolysis from the promiscuous activity of 1-phosphofructokinase (FruK). We evolved a hexose-degradation-deficient *Escherichia coli* strain on glucose and recovered non-synonymous mutations in *fruK*. Reverse engineering, gene deletions and transcript quantification identified a modified and strongly overexpressed *fruK* as the determinant of the growth phenotype. Purified FruK variants phosphorylated fructose-6-phosphate *in vitro*, the best variant with ∼170-fold improved catalytic efficiency over the wildtype enzyme. Crystal structures at up to 1.4 Å resolution captured FruK with ADP and either F1P, F6P or FBP bound. They identify a single active-site lysine that excludes the alternative substrate, and which *E. coli* FruK retains despite forgoing a 26 % gain in efficiency on its own native reaction. Since F1P, the product of the reverse reaction of FruK, is the effector of the global regulator Cra, we propose that this residue keeps glycolytic flux separate from transcriptional control. The same active-site motifs occur naturally across FruK clades, and a hexose-degradation-deficient *Corynebacterium glutamicum* strain converged on the same adaptation, indicating a common strategy across the Gram divide. The committed step of EMP glycolysis therefore lies a single substitution away within a widespread sugar kinase family, where it appears to be actively suppressed.

## Introduction

Darwinian evolution is driven by genetic variation and selection. Organisms with traits that improve fitness are more likely to pass on those traits, while less beneficial traits fade away with some exceptions. At the metabolic level, natural selection typically favors enzyme variants that catalyze reactions more efficiently, *i.e.* higher flux per enzyme, gradually optimizing biochemical pathways over time^1^. Active site promiscuity plays an important role in these evolutionary processes. The ability of enzymes to catalyze different reactions or act on various substrates^2,3^, allows them to quickly diversify, adapt new functions and fill new niches^4^. Hence, enzyme promiscuity is a key driver of evolutionary innovation and phenotypic plasticity^5,6^.

Glycolysis is one of the most important and dominant metabolic pathways in nature. Multiple glycolytic pathways have been found across all domains of life^7^. *E. coli* natively employs a phosphotransferase system (and can evolve to use a permease/glucokinase pair) to import and phosphorylate glucose^8^ into glucose-6-phosphate (G6P). G6P is then degraded via three different glycolytic routes: The Embden-Meyerhof-Parnas pathway, the Entner-Doudoroff pathway and the oxidative pentose phosphate pathway (Fig. 1, Supplementary Fig. 1). Previous work of Miller and Raines provided compelling experimental evidence that enzyme promiscuity allows *E. coli* to adapt to a loss of phosphorylation-driven glucose import ^9,10^. By constructing an *E. coli* strain deficient in both its phosphotransferase system and glucokinase (*glk*), they demonstrated that at least four ROK-family sugar kinases, namely NanK, YajF, YcfX, and AlsK harbor latent glucokinase activities and are sufficient to rescue growth of the auxotrophic strain on glucose minimal medium.

**Figure 1:**
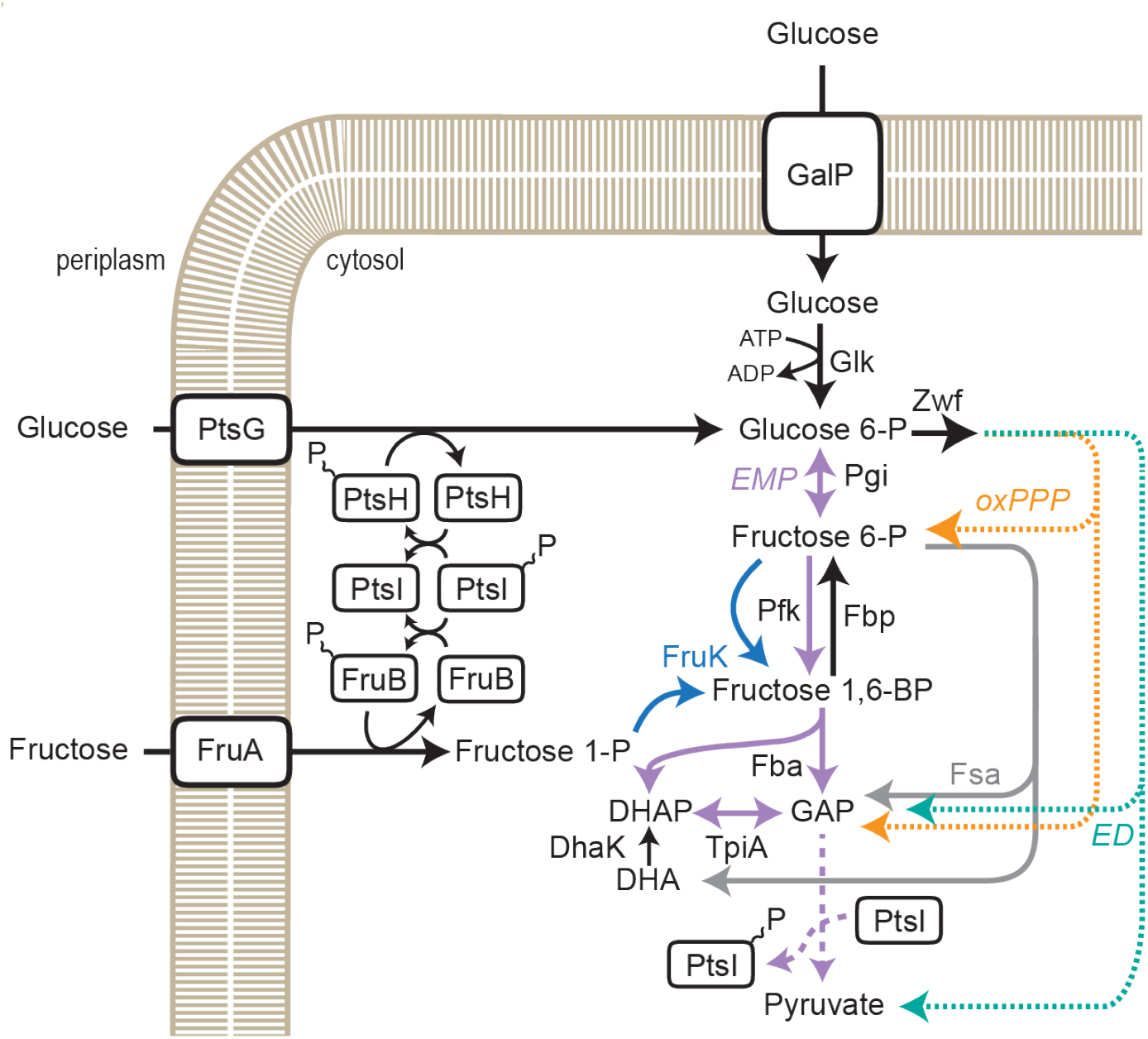
Main glucose and fructose degradation routes in *E. coli*. Glucose and fructose import and degradation to pyruvate are shown. Glucose and fructose both have a phosphoenolpyruvate-dependent phosphotransferase system (PTS; PtsG and FruA, supplied with phosphoryl groups through PtsH and FruB respectively, which in turn are both phosphorylated by enzyme I (PtsI)) and alternative import and activation options (GalP/Glk; only the canonical of 4 potential utilization routes is shown for fructose*^12,13^*, Supplementary Fig. 1). Main glycolytic routes are labeled and color-coded, Embden-Meyerhof-Parnas (EMP) pathway in violet, Entner-Doudoroff (ED) pathway in turquoise, oxidative pentose phosphate pathway (oxPPP) in yellow and fructose-6-phosphate aldolase (Fsa) route in grey. The canonical and novel reactions of 1-phosphofructokinase (FruK) are shown in blue. Phosphoglucose isomerase (Pgi), Fructose bisphosphate phosphatase (Fbp), 6-phosphofructokinase (Pfk), Fructose bisphosphate aldolase (Fba), Triosephosphate isomerase (TpiA), Dihydroxyacetone kinase (DhaK), Dihydroxyacetone phosphate (DHAP), Glyceraldehyde-3-phosphate (GAP).

Building on this framework, we sought to explore whether analogous recruitment events could support the emergence of novel glycolytic routes by introducing targeted deletions in the Embden–Meyerhof–Parnas, the Entner–Doudoroff and the oxidative pentose phosphate pathways, forcing the cell to evolve alternative means of glucose catabolism. To this end, we systematically abolished all known glucose degradation routes in *Escherichia coli*, by deletion of 6-phosphofructokinase (*pfkAB*), glucose-6-phosphate dehydrogenase (*zwf*) and fructose-6-phosphate aldolase (*fsaAB*)^11^. We then evolved this hexose-degradation-deficient strain on glucose via adaptive laboratory evolution. Suppressor mutants were isolated and were found to harbor amino acid substitutions in the active site of 1-phosphofructokinase (FruK). We employed genetic and enzymatic approaches to biochemically characterize several FruK variants and showed that all were able to perform the 6-phosphofructokinase reaction with high catalytic efficiency, readily bypassing the deletion of *pfkAB*. We then solved the crystal structures of FruK from *E. coli* and several of its mutant variants with bound substrates and products. Finally, we found the same FruK based bypass in *C. glutamicum*, show that its FruK natively harbors our mutated active site motif and correspondingly catalyzed the 6-phosphofructokinase reaction *in vitro.* A single active-site residue therefore separates the two phosphofructokinase reactions, and its substitution is sufficient to re-establish EMP glycolysis, suggesting that the committed step of the pathway lies within reach of a promiscuous sugar kinase scaffold.

## Results

### *E. coli* strains lacking all known hexose degradation routes evolve to grow on glucose

To uncover potentially novel glycolytic routes, we first set out to delete all known hexose degradation pathways*. E. coli* strains deleted in both 6-phosphofructokinases (*pfkA* and *pfkB*) are unable to grow on glucose^14^, due to NADPH accumulation from the Entner-Doudoroff and oxidative pentose phosphate pathway, respectively^15^. Having in mind that soluble transhydrogenase can rebalance NAD(P)H pools^16^, we additionally deleted the *pfkA*, *pfkB* and *zwf* genes encoding 6-phosphofructokinase and glucose-6-phosphate dehydrogenase to fully abolish growth on glucose. This Δ*pfk* Δ*zwf* deletion strain (PZ strain) was indeed unable to grow on glucose. Nevertheless, after about 5 days of extended cultivation growth on glucose minimal media re-occurred (Supplementary Fig. 2A). When isolates were re-inoculated into glucose minimal media, they displayed immediate growth, confirming the emergence of (stable) suppressor mutants. We speculated that these suppressors involved fructose-6-phosphate aldolase, which is encoded by two *E. coli* genes (*fsaA* and *fsaB*) and was recently shown to circumvent *pfk* deletions^11^. To test if fructose-6-phosphate aldolase caused the growth restoration we then also deleted both *fsa* genes resulting in strain Δ*pfk* Δ*zwf* Δ*fsaA* Δ*fsaB* (PZF strain). However, strain PZF was still able to grow on glucose after several days (Supplementary Fig. 2B), indicating an alternative bypass mechanism.

### Mutations in 1-phosphofructokinase (FruK) allow growth on glucose

To understand the cause of the suppressor mutants, we sequenced the genomes of ten clonal isolates of PZ and PZF mutants and mapped all genetic differences relative to their parents, *i.e.*, the un-mutated PZ and PZF strains, which contained a *cra* and *napD* mutation compared to *E. coli* SIJ488 wildtype (Tab. 1). Notably, in eight of ten suppressor clones, the gene *fruK* encoding 1-phosphofructokinase was affected by non-synonymous mutations. In seven cases, lysine 95 was mutated into histidine, threonine or asparagine (PZF^95H/T/N^), while in one case leucine 265 was changed into a phenylalanine (PZF^265F^). These different FruK mutations translated into distinct growth phenotypes. Variants PZF^95H^, PZF^95T^ and PZF^95N^ reproducibly grew on glucose with 2, 3 and 4 hours doubling time, respectively, while PZF^265F^ showed more than eleven hours of doubling time (Fig. 2). In two independent isolates we also found mutations in *apaH* and *ptsI* encoding diadenosine tetraphosphatase and PTS enzyme I, respectively. However, these mutants showed an extended lag phase on glucose and were therefore not further pursued in this study. To dissect the observed mutations and explain the glucose growth phenotype, we instead focused on the *fruK* mutations. During growth on fructose FruK catalyzes the ATP-dependent phosphorylation of fructose-1-phosphate (F1P; derived from the dedicated fructose PTS) into the glycolytic intermediate fructose-1,6-bisphosphate (FBP) (Fig. 1)^17^. While we initially entertained several scenarios of FruK induced metabolic bypasses (Supplementary Fig. 3), the evolution of similar mutants on D-xylose and D-ribose led us to speculate that FruK was directly phosphorylating fructose-6-phosphate (F6P) to FBP (Supplementary Fig. 2A, B).

**Figure 2:**
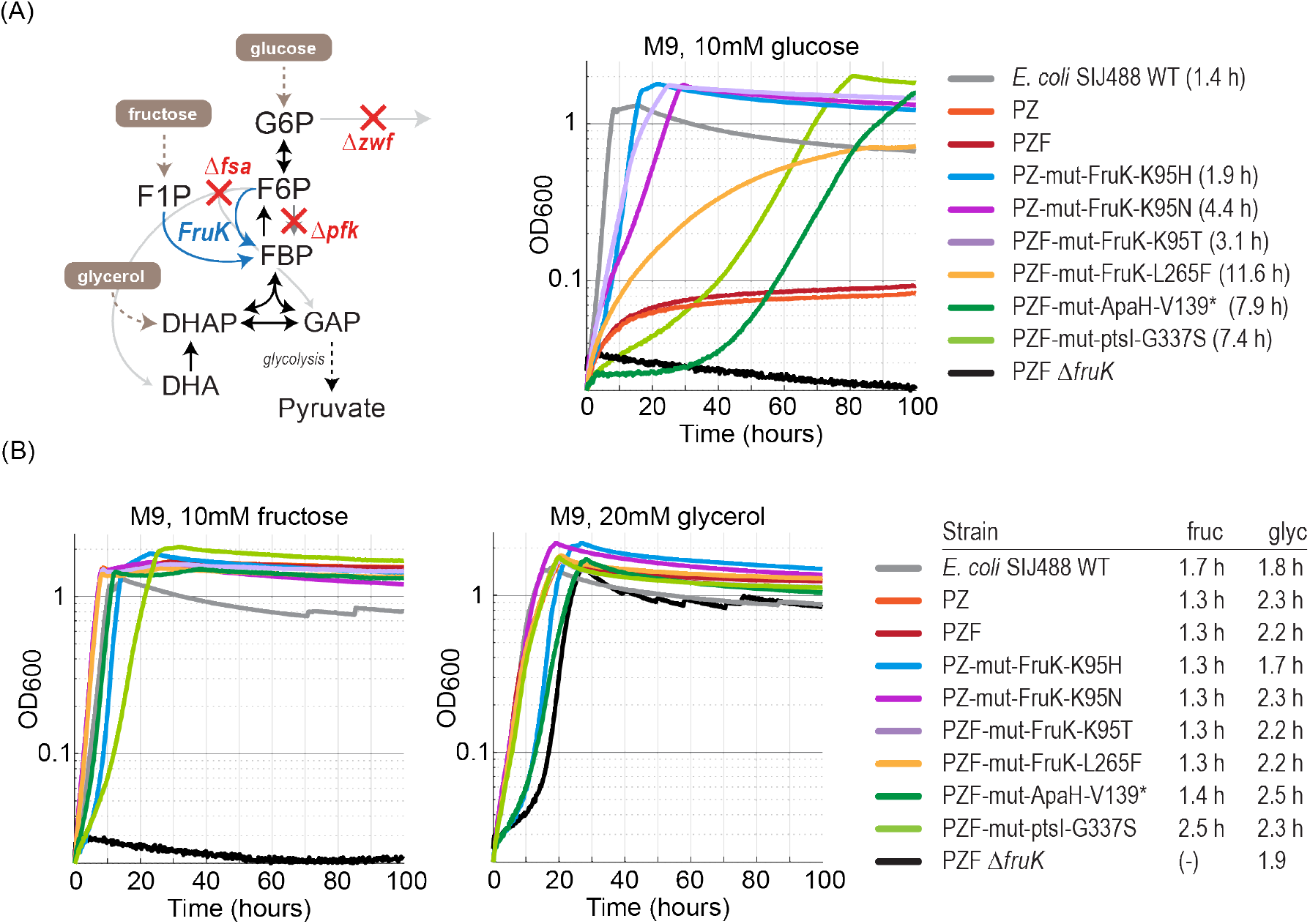
Growth of *E. coli* FruK mutants on glucose. A, Simplified view of sugar-phosphate metabolism and growth profiles of PZ and PZF deletion strains as well as with and without fruK mutations grown in M9 media with either glucose (A), fructose (B) or glycerol (C). DHA, dihydroxyacetone; DHAP, dihydroxyacetone-phosphate; F1P, fructose-1-phosphate; F6P, fructose-6-phosphate; FBP, fructose-1,6-bisphosphate; *fsa*, fructose-6-phosphate aldolase; *fruk*, 1-phosphofructokinase; G6P, glucose-6-phosphate; GAP, glyceraldehyde-3-phosphate; PEP, phosphoenolpyruvate; *pfk*, 6-phosphofructokinase; PYR, pyruvate; *zwf*, glucose-6-phosphate dehydrogenase.

**Table 1:** Whole genome sequencing results of PZ and PZF parental strains and the derived glucose growth mutants. Parental strains are labeled PZ and PZF respectively, PZ mutants 1-4 (are labeled 1-4) and PZF mutants 1-6 were named (5-10). Mutations are given relative to the SIJ488 reference genome (GenBank: CP132594).

| **Gene** | **Identifier** | **Position** | **Mutation** | **PZ** | **PZF** | **1** | **2** | **3** | **4** | **5** | **6** | **7** | **8** | **9** | **10** |
|---|---|---|---|---|---|---|---|---|---|---|---|---|---|---|---|
| *cra* | ECK0081 | 88,283 | Q86*<br>(CAG→TAG) | x | x | x | x | x | x | x | x | x | x | x | x |
| *napD* | ECK2199 | 2,302,485 | Q58Q<br>(CAG→CAA) | x | x | x | x | x | x | x | x | x | x | x | x |
| *gltP* | ECK4070 | 4,304,457 | Intergenic<br>(+587/+55) | x | x | x | x | x | x | x | x | x | x |  | x |
| *fruK* | ECK2161 | 2,261,726 | K95H<br>(AAG→CAT) |  |  | x | x |  |  |  |  |  |  |  |  |
| *fruK* | ECK2161 | 2,261,728 | K95H<br>(AAG→CAT) |  |  | x | x |  |  |  |  |  |  |  |  |
| *fruK* | ECK2161 | 2,261,726 | K95N<br>(AAG→AAT) |  |  |  |  | x | x |  |  |  |  |  |  |
| *fruK* | ECK2161 | 2,261,727 | K95T<br>(AAG→ACG) |  |  |  |  |  |  | x | x | x |  |  |  |
| *fruK* | ECK2161 | 2,261,216 | L265F<br>(TTG→TTT) |  |  |  |  |  |  |  |  |  | x |  |  |
| *apaH* | ECK0050 | 50,839 | V139*<br>(Δ1 bp at 384nt) |  |  |  |  |  |  |  |  |  |  | x |  |
| *ptsI* | ECK2411 | 2,534,719 | G337S<br>(GGC→AGC) |  |  |  |  |  |  |  |  |  |  |  | x |

### Genetic approaches to analyze FruK’s contributions to growth phenotypes

To confirm that the mutations in *fruK* were the reason for the restored growth of the mutant isolates we followed a two-pronged approach. First, we deleted *fruK* in the fast-growing mutants PZ^95H^ and PZF^95T^. Deletion of *fruK* resulted in unhealthy *E. coli* strains, which adapted over the course of several re-streaks on LB plates and were accompanied by a truncation of Pts enzyme 1 after glutamate 422^18^. Adapted strains PZ^95H^Δ*fruK* and PZF^95T^Δ*fruK* (named PZFK) were unable to grow on glucose, even upon extended cultivation (Fig. 2A, data for PZF^95T^Δ*fruK* shown as black line) and on xylose or ribose (Supplementary Fig. 2C), supporting the role of FruK in growth rescue. Since these strains also carry a truncation of PTS enzyme 1, we addressed a possible contribution of impaired glucose import by plasmid-borne complementation of PZFK* (see below).

Second, we reverse engineered the identified FruK mutations in the ancestor PZF strain, using multiplex automated genome engineering (MAGE)^19^. We introduced the four FruK substitutions K95H, K95T, K95N and L265F individually into the PZF strain and directly selected for growth on glucose minimal media plates after one MAGE round (Supplementary Fig. 4). While we could obtain MAGE clones for K95H and K95T, we did not recover L265F, and K95N only in combination with additional mutations (e.g., 3 bp deletions, Fig. 3A, C). Sequencing more clones, we identified additional mutations in FruK, such as K95S, G110A and G110S that directly enabled growth (Fig. 3A, C), and mutations that synergistically improved growth (e.g., G110S, Q84R or T34I in combination with K95T). Taken together these experiments confirmed that a modified *fruK* gene is crucial to metabolize glucose in a strain devoid of 6-phosphofructokinase and glucose-6-phosphate dehydrogenase activity.

**Figure 3:**
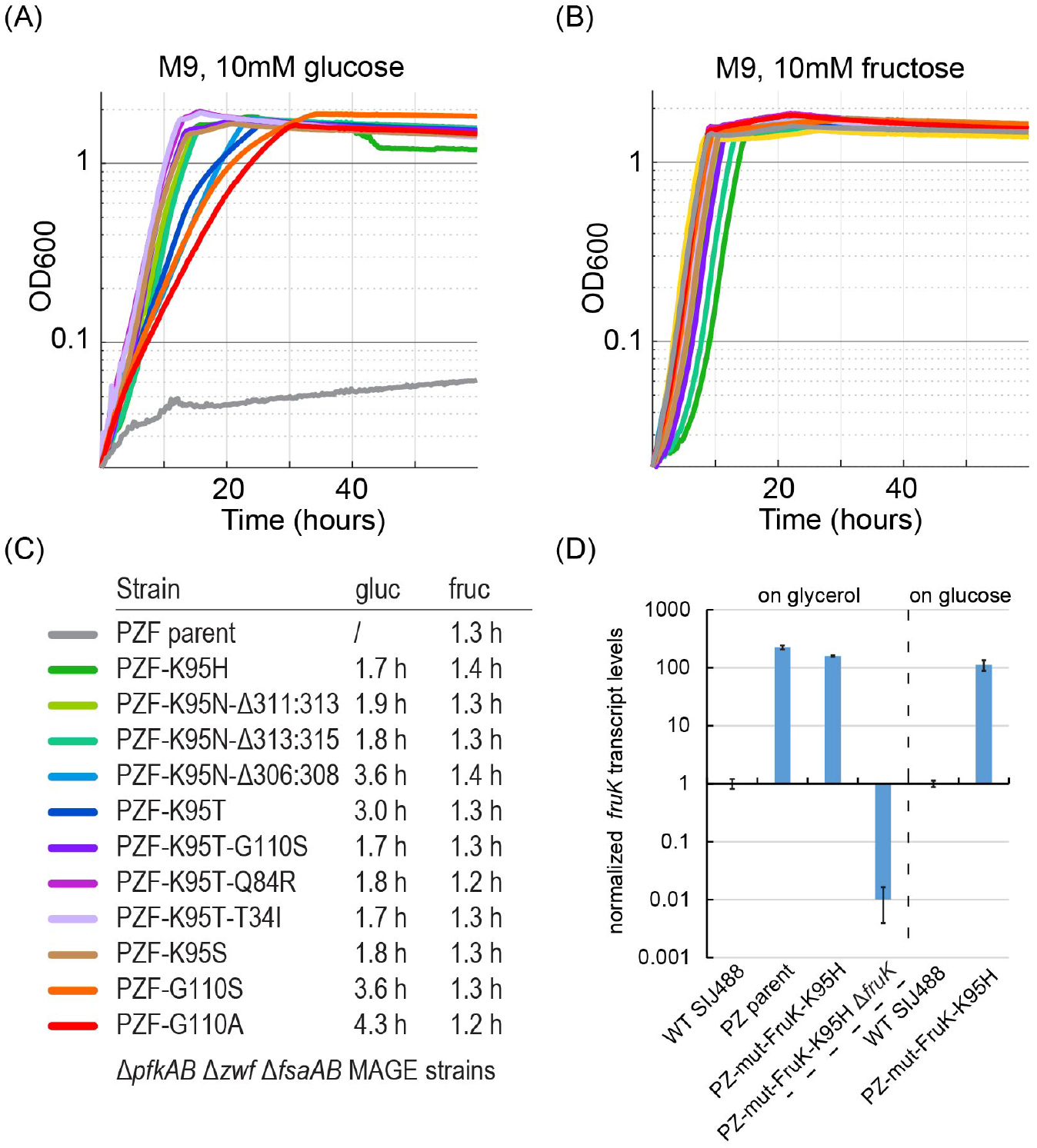
Reverse-engineered fruK mutations using MAGE restore growth on glucose in a fruK-overexpressing background. **A-C**, *E. coli* PZF FruK-MAGE strains were cultivated in glucose (gluc) or fructose (fruc) M9 minimal media. While the FruK substitutions K95H and K95T could be reverse engineered several additional substitutions especially accompanying K95N were observed. PZF-K95N-Δ311:313, -Δ313:315 and - Δ306:308 indicate 3 different deletions of 3 bps on top of the FruK-K95N substitution. L265F did not re-occur using MAGE. **D**, Quantification of FruK transcript levels in *E. coli* PZ and PZ mutant strains relative to *E. coli* WT as measured by quantitative reverse transcriptase PCR. The transcript levels of *fruK* were upregulated by two orders of magnitude. Error bars show standard deviations. FruK transcript abundance of the delta *fruK* strain was indistinguishable from the water control.

To investigate whether these mutations compromised the original function of *fruK* in fructose metabolism *i.e*., the phosphorylation of F1P into FBP, we grew the PZF mutant strains on fructose minimal media (Fig. 2B and 3B, C). Notably, we did not observe any significant differences in growth on fructose minimal medium compared to the parental and wildtype strains (Fig. 2B and 3B, C). Thus, our experiments suggested that all FruK variants maintained their initial F1P phosphorylation activity but additionally acquired a new function, possibly a F6P phosphorylation activity, that allowed these variants to also operate in glucose utilization.

To confirm production of *fruK* on glucose as carbon source, we determined *fruK* expression levels with quantitative reverse transcriptase PCR (qRT-PCR) (Fig. 3D, Supplementary Fig. 5). Our analysis revealed that *fruK* was upregulated in the parental and mutant strains by more than two orders of magnitude when compared to *E. coli* SIJ488 wildtype grown on glucose. This was likely caused by truncation of Cra in our parental strains (see above, Tab. 1), which was formerly known as FruR and directly controls FruK expression^20^. In summary, this data suggested that mutation of FruK in combination with *fruK* overexpression contributes to the suppressor phenotype, which we speculated to be based on a novel F6P phosphorylation activity (Supplementary Fig. 3).

### Biochemical characterization identifies latent 6-phosphofructokinase activity within FruK variants

Native FruK catalyzes the ATP-dependent phosphorylation of F1P into FBP (Fig. 1)^17^, while Pfk canonically phosphorylates F6P at position 1. Although FruK and PfkB are both members of the ribokinase enzyme family^21^, they share only about 30 % of amino acid identity^22^. To test whether FruK is also able to catalyze the phosphorylation of F6P, we purified His-tagged versions of *E. coli* wildtype FruK, and several mutant variants and performed a classical pyruvate kinase/lactate dehydrogenase coupled assay. Indeed, while *E. coli* FruK showed a *k*_cat_ of ∼200 sec^-1^ at a *K*_M_ of ∼0.2 mM for its native substrate F1P, it also displayed a basal activity of less than 1% with F6P, that is a *k*_cat_ of ∼1 sec^-1^ at a *K*_M_ of 3.5 mM (Tab. 2, Supplementary Fig. 6). Interestingly, FruK variant K95H exhibited an almost 170-fold increase in catalytic efficiency with F6P compared to wildtype (∼10-fold improved *k_cat_* and 16-fold decreased *K*_M_), without sacrificing catalytic efficiency for its native substrate F1P (Tab. 2, Supplementary Fig. 6). All other FruK mutations (K95N, K95T, K95T/G110S and K95T/T34I) mainly increased the *k*_cat_ for the F6P reaction between ∼5- to 10-fold, while strongly decreasing catalytic efficiency for the native reaction (by ∼10-fold). We also tested activity of our FruK variants on unphosphorylated fructose, but observed no activity, indicating that one phosphoryl group on the substrate was crucial for substrate binding and/or catalysis.

**Table 2:** Comparison of biochemical properties of different *E. coli* FruK variants.

|  | *Escherichia coli* |  |  |  |  |  |
|---|---|---|---|---|---|---|
| Variant | WT | K95H | K95T | K95N | K95T/G110S | K95T/T34I |
|---|---|---|---|---|---|---|
|  | Fructose-1-phosphate |  |  |  |  |  |
| *k*<sub>cat</sub> (s<sup>-1</sup>) | 206 | 102 | 180 | 128 | 131 | 182 |
| app. *K*<sub>M</sub> (µM) | 182 | 72 | 1850 | 1860 | 576 | 540 |
| *k*<sub>cat</sub>/*K*<sub>M</sub> (M<sup>-1</sup> × s<sup>-1</sup>) | 1.1x10<sup>6</sup> | 1.4x10<sup>6</sup> | 9.7x10<sup>4</sup> | 6.9x10<sup>4</sup> | 2.3x10<sup>5</sup> | 3.4x10<sup>5</sup> |
|  | Fructose-6-phosphate |  |  |  |  |  |
| *k*<sub>cat</sub> (s<sup>-1</sup>) | 1.4 | 14.0 | 6.8 | 6.2 | 12.3 | 15.7 |
| app. *K*<sub>M</sub> (µM) | 3490 | 213 | 4180 | 8890 | 1660 | 2100 |
| *k*<sub>cat</sub>/*K*<sub>M</sub> (M<sup>-1</sup> × s<sup>-1</sup>) | 4x10<sup>2</sup> | 6.6x10<sup>4</sup> | 1.6x10<sup>3</sup> | 7.0x10<sup>2</sup> | 7.4x10<sup>3</sup> | 7.5x10<sup>3</sup> |

### Structural investigation of *E. coli* FruK variants reveals molecular basis for the 6-phosphofructokinase activity

To investigate structural changes facilitating the shift in substrate specificity and rate of catalysis, we first co-crystallized *E. coli* FruK wildtype, as well as variants K95H and K95T/G110S with ADP, Mg^2+^, and F1P (Fig. 4, Supplementary Fig. 7). We obtained high resolution structures (between 1.41 and 1.55 Å) with the native substrate bound to the active site (Fig. 4). However, much to our surprise, despite repeated attempts to co-crystallize these variants with F6P instead, all datasets again revealed F1P bound at the active site, indicating that FruK had converted F6P into F1P.

**Figure 4.**
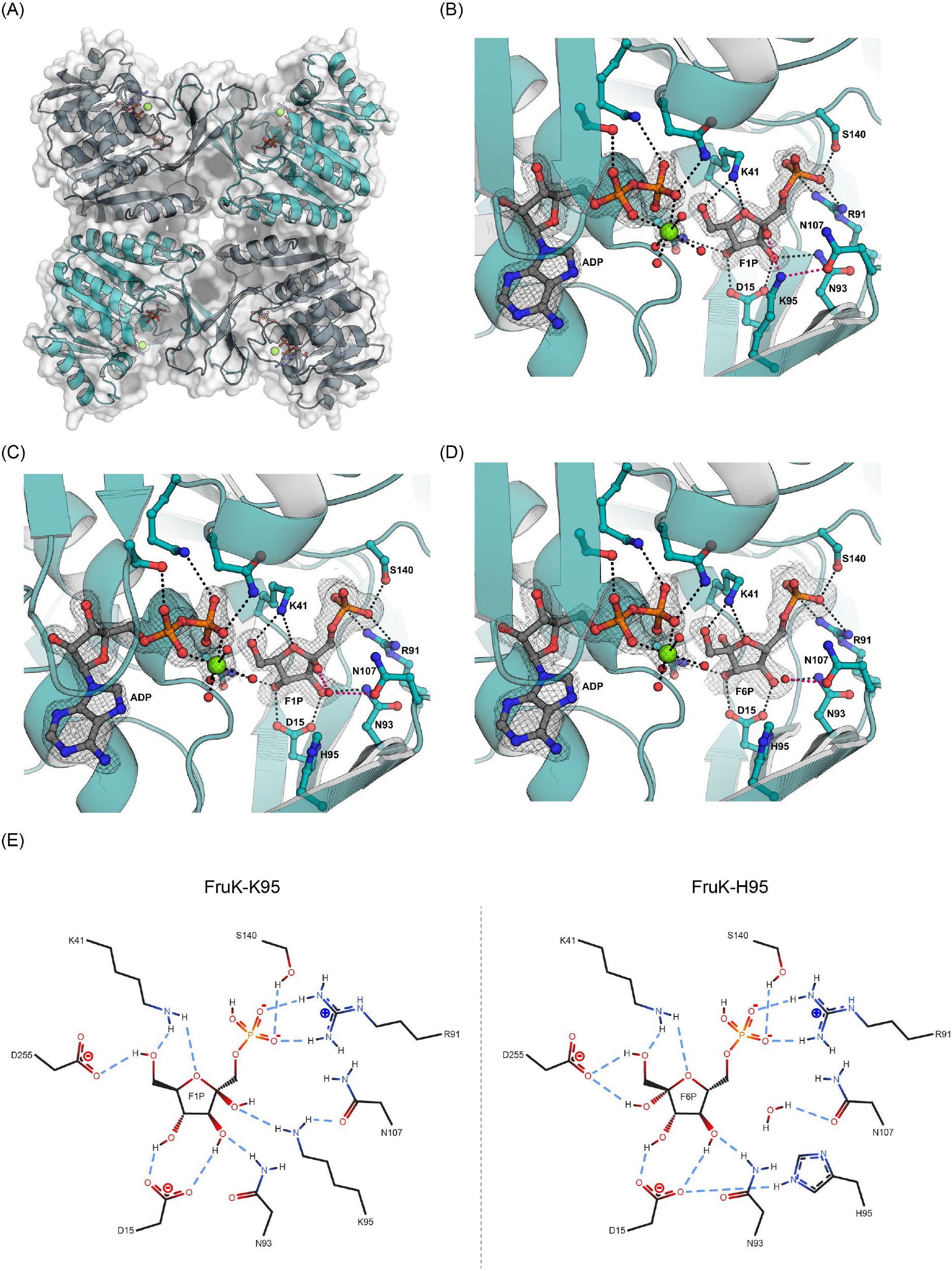
Crystal structure of the tetrameric assembly of the *E. coli* FruK variants and proposed mechanism. **A**, shown is a cartoon representation of FruK. The different subunits are colored alternating in light and dark teal. A semi-transparent surface representation is superposed. Bound substrates (ADP, F1P, and Mg^2+^) are shown as balls and sticks to highlight the locations of the respective active sites. **B**, active site of the WT variant of the *E. coli* FruK with bound substrates. Polar interactions between sidechains and ligands are depicted by dashed lines. Interaction with the residue K95 are highlighted in pink, as this position was found to be substituted in most of the mutant variants. A simulated annealing omit map is shown at a σ of 4.0 around the ligands. **C** and **D** active site of the K95H variant of the *E. coli* FruK with bound F1P and F6P, respectively. Polar interactions between sidechains and ligands are depicted by dashed lines. Interactions with a water molecule occupying the space of the substituted residue K95 (here H95) are highlighted in pink. Simulated annealing omit maps are shown at a σ of 4.0 for ADP, at σ of 2.0 for F1P, and σ of 1.5 for F6P. **E**, Scheme depicting the modes of binding for F1P in the wild type enzyme and F6P in case of the K95H variant.

We investigated the apparent “mutase” activity in more detail, using LC/MS (Supplementary Fig. 8). F6P to F1P conversion only occurred when we used ADP as a co-substrate but not when ATP was present in saturating or excess amounts. Based on these findings, we speculated that the ADP used during co-crystallization contained traces of ATP in catalytic amounts, so that FruK catalyzed the forward reaction, i.e. formation of FBP from F6P, and — favored by the high ADP concentrations— subsequently the backreaction resulting in F1P accumulation, which preferentially binds at the active site, over time (Supplementary Fig. 8). We therefore attempted soaking the crystals with excess F6P immediately before freezing, which indeed allowed us to obtain one structure for variant K95H with bound F6P, the initial substrate, at a resolution of 1.85 Å (Fig. 4D) and one structure of variant K95T/G110S with the product FBP at 1.55 Å (Supplementary Fig. 7), essentially trapping all substrates and products of the forward and backward reactions, respectively.

Notably, the structural comparison of FruK wildtype with mutant variants provided an explanation for the K95 mutation (Fig. 4E). In the wildtype enzyme, this lysine residue engages in a hydrogen bond (2.8 Å) with the hydroxyl group at the C2 of the fructofuranose ring (Fig. 4B). This strongly favors F1P as substrate and even prevents binding of F6P, for which FruK needs to bind the furanose ring in a “flipped” orientation to enable C1 phosphorylation. Substitution of K95 with shorter residues (*i.e.* histidine, threonine) creates space for another water molecule (Fig. 4C, D, E). This water molecule is less fixed in space than the original ε-amino group of K95, now allowing both, F1P and F6P binding (Fig. 4C). Unfortunately, our crystal structures did not provide direct insights into the effects of the G110S or T34I substitutions, which might be rather based on changes in rigidity and/or sub-Ångstrom positioning shifts of secondary structure elements. In summary, these structural investigations allowed us to visualize individual steps along the reaction coordinate and provided insights into the molecular basis of the newly acquired F6P phosphorylation activity of the most relevant FruK variants.

### Promiscuous 6-phosphofructokinase FruK activity occurs throughout broad enzyme families

Having identified molecular determinants for F6P phosphorylation activity, we next sought to investigate the natural diversity of microbial FruK homologs. Filtering 5,000 uniprot sequences, we performed a phylogenetic analysis of the sequence space, which showed that FruK separated into different subclades (Fig. 5). Notably, these subclades displayed diversity at residue 95. One prominent example was the subclade including FruK1 of the gram-positive bacterium *Corynebacterium glutamicum*. In this subclade lysine 95 was replaced by a threonine, indicating that these enzymes might possess some native F6P phosphorylation activity. Taking our previous findings together with a previous literature report on FruK mutations in glucose evolved *C. glutamicum* strains^23^, we aimed to investigate whether *C. glutamicum* FruK also possesses 6-phosphofructokinase activity.

**Figure 5:**
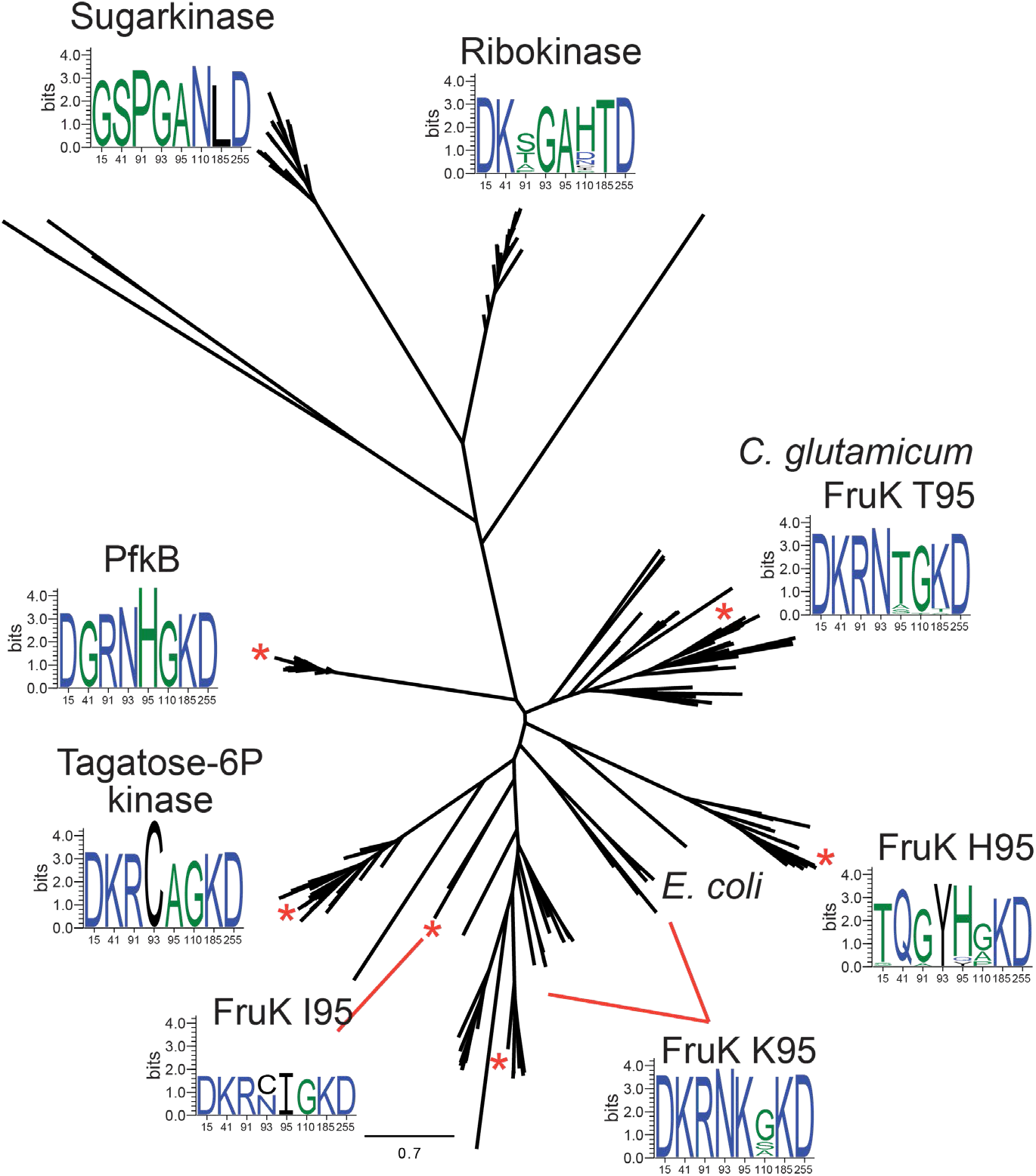
Conservation of the FruK active-site region across sugar kinase families. Unrooted maximum-likelihood phylogeny of FruK (1-phosphofructokinase), ribokinase, sugar kinase, tagatose-6-phosphate kinase, and PfkB (6-phosphofructokinase-2) homologs, with pyridoxal kinase (PdxK) as the outgroup. *E. coli* FruK and chosen representative homologs are indicated on the tree (explicitly or by an asterisk). For each clade, a sequence logo shows amino acid conservation at eight active-site positions (numbered according to *E. coli* FruK). FruK homologs partition into sub-clades with varying residues at active site position 95. Colors in sequence logos represent similarities in size, charge and polarity, scale bar represents the number of substitutions per site.

Analogous to our efforts with *E. coli* (see above), we generated a *C. glutamicum* Δ*pfk* Δ*zwf* double knockout strain that cannot metabolize F6P via EMP glycolysis or the pentose phosphate pathway. This strain was able to grow on fructose but not glucose as the sole carbon source (Fig. 6). However, prolonged incubation on glucose resulted in suppressor mutants, of which we analyzed four clones in more detail. All four clones showed similar growth behavior on glucose minimal medium and contained one of two different mutations in the *fruK1* gene: a substitution L29F or G104A, and three out of four showed an additional mutation in transcriptional regulator SugR (Cgl1931 Cg2115, Tab. 3). To confirm that growth on glucose was related to FruK1 (Cgl1935 (cg2119)), we generated strain Δ*pfk* Δ*zwf* Δ*fruK1*, which did not show growth on glucose minimal medium even after prolonged incubation for > 12 days.

**Figure 6:**
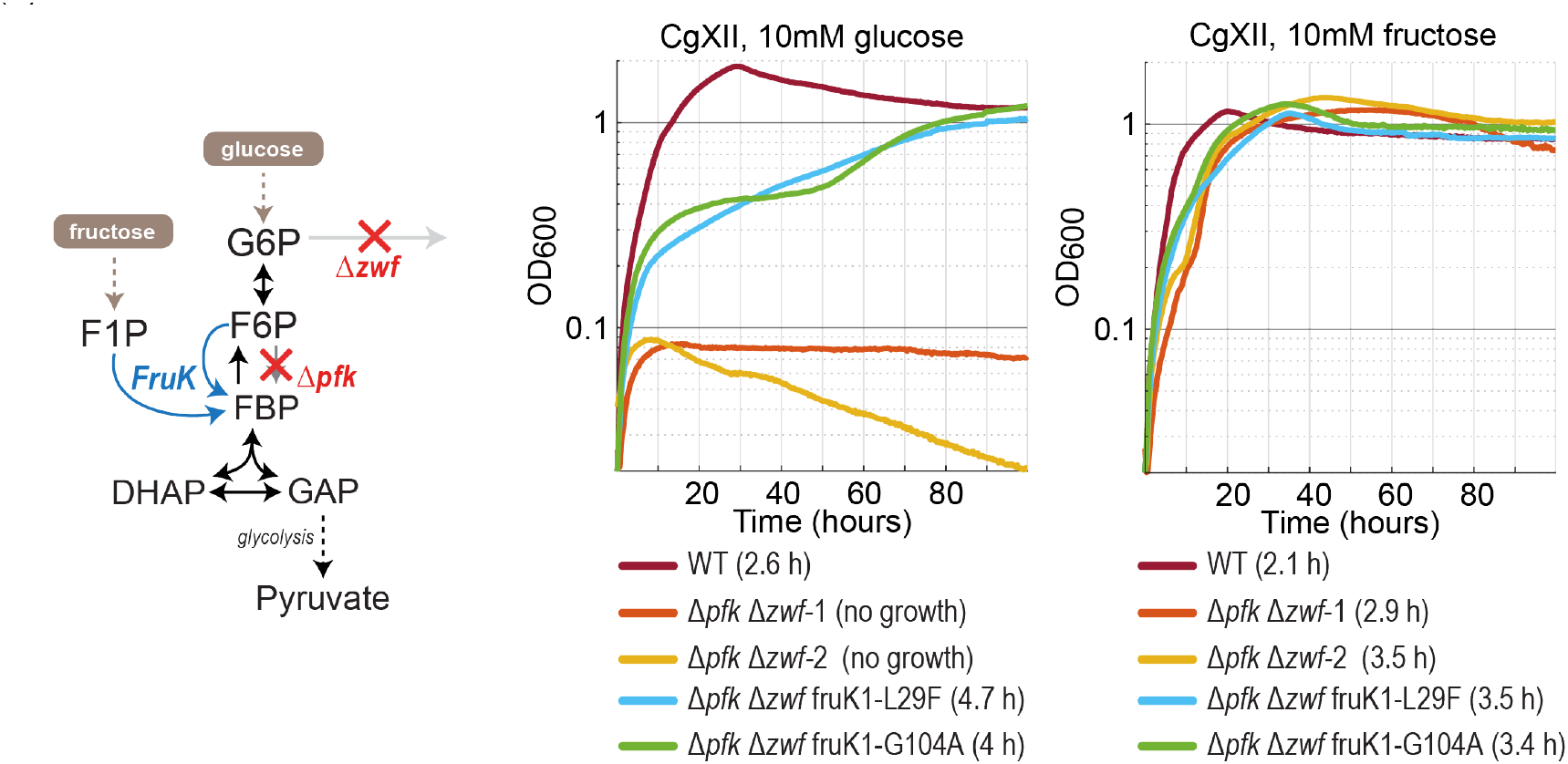
Growth of *C. glutamicum* Δ*pfk* Δ*zwf* and derived mutant isolates on glucose or fructose. Strains were grown in CgXII media supplemented with 10 mM glucose or fructose in a 96-well plate reader. Identified FruK mutations are indicated as well as the corresponding doubling times of the strains. DHAP, dihydroxyacetone-phosphate; F1P, fructose-1-phosphate; F6P, fructose-6-phosphate; FBP, fructose-1,6-bisphosphate; *Fruk*, 1-phosphofructokinase; G6P, glucose-6-phosphate; GAP, glyceraldehyde-3-phosphate; *pfk*, 6-phosphofructokinase; PYR, pyruvate; *zwf*, glucose-6-phosphate dehydrogenase.

**Table 3:** Mutations found in the glucose grown C. glutamicum Δ*pfk* Δ*zwf* strains relative to the parental strain.

| **Gene** | **Identifier 1** | **Mutation** | **PZ mut 1** | **PZ mut 2** | **PZ mut 3** | **PZ mut 4** |
|---|---|---|---|---|---|---|
| FruK1 | Cg2119 | L29F | x | x | x |  |
| FruK1 | Cg2119 | G104A |  |  |  | x |
| SugR | Cg2115 | G55D |  | x |  |  |
| SugR | Cg2115 | lexA insertion |  |  | x |  |
| SugR | Cg2115 | tnp insertion |  |  |  | x |

We also transferred *C. glutamicum fruK1* wildtype and the different variants to the *E. coli* PZ, the PZF and the PZFK* strains and tested for growth complementation (Fig. 7, Supplementary Fig. 9 and 10). The PZFK* strain was created by re-evolving growth of the Δ*pfk* Δ*zwf* Δ*fsaAB* Δ*fruK* strain on glucose while overexpressing FruK to address the import issue caused by the truncated PTS enzyme 1. This way we were able to untangle whether a potential “no growth” phenotype of the PZFK strain was due to a missing PFK-like reaction or due to missing glucose import. Notably, all *C. glutamicum* FruK1 variants, including the wildtype enzyme complemented growth of all strains, including strain PZFK* with truncated PTS enzyme 1 on glucose minimal medium when overexpressed from the pZ vector system, indicating that these enzymes possessed F6P phosphorylation activity (Fig. 7A). Moreover, these complementations also restored growth of the PZF strain in xylose minimal medium (Fig. 7B), where xylose generates stoichiometric amounts of F6P (i.e., 2:1 F6P:glyceraldehyde-3-phosphate) that accumulates intracellularly, inhibiting growth without functional FruK. Growth on fructose remained largely unchanged except for the PZFK* strain, where the lack of the truncated PTS enzyme 1 abolished fructose import, as expected. Overall, these results demonstrated that *C. glutamicum* FruK1 (and variants) allowed for glucose utilization.

**Figure 7:**
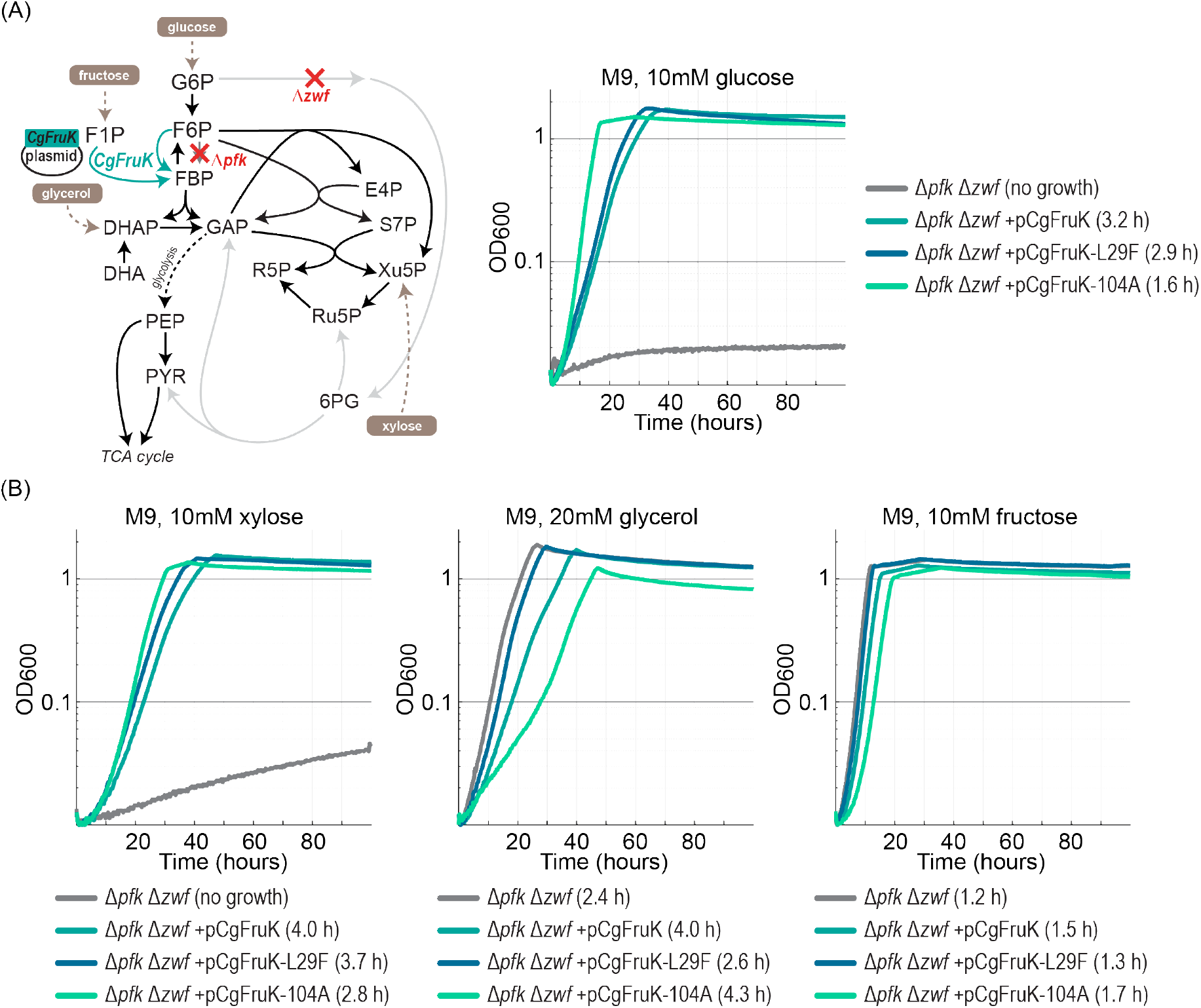
Plasmid complementation experiment of the *E. coli* Δ*pfk* Δ*zwf* strain with *C. glutamicum* FruK1 variants restore growth on glucose and xylose. Strains were cultivated in M9 media supplemented with either 10mM glucose, 10mM xylose, 20mM glycerol or 10mM fructose.

Biochemical characterization confirmed that FruK1 wildtype already contained considerable activity with F6P (k_cat_ ∼5 sec^-1^ at a *K*_M_ of ∼350 µM) compared to F1P (kcat 20 sec^-1^, at a *K*_M_ of ∼20 µM, Tab. 4, Supplementary Fig. 6G-I). Variant L29F behaved very similar to the wildtype, while FruK 1 G104A showed an increase of ∼70% in *k*_cat_ for the novel substrate F6P (∼10 sec^-1^) but also almost one order of magnitude improved activity on the natural substrate, F1P (∼140 sec^-1^). Thus, similarly to *E. coli* FruK, FruK1 can be further improved for its native activity and F6P phosphorylation through a single mutation.

**Table 4:** Comparison of biochemical properties of different *C. glutamicum* FruK variants.

|  | *Corynebacterium glutamicum* |  |  |
|---|---|---|---|
| Variant | WT | L29F | G104A |
|  | Fructose-1-phosphate |  |  |
| *k*<sub>cat</sub> (s<sup>-1</sup>) | 20 | 25 | 141 |
| app. *K*<sub>M</sub> (µM) | 19 | 21 | 31 |
| *k*<sub>cat</sub>/*K*<sub>M</sub> (M<sup>-1</sup> × s<sup>-1</sup>) | 1.1x10<sup>6</sup> | 1.2x10<sup>6</sup> | 4.6x10<sup>6</sup> |
|  | Fructose-6-phosphate |  |  |
| *k*<sub>cat</sub> (s<sup>-1</sup>) | 5.8 | 6.4 | 10.1 |
| app. *K*<sub>M</sub> (µM) | 335 | 540 | 940 |
| *k*<sub>cat</sub>/*K*<sub>M</sub> (M<sup>-1</sup> × s<sup>-1</sup>) | 1.7x10<sup>4</sup> | 1.2x10<sup>4</sup> | 1.1x10<sup>4</sup> |

## Discussion

In this study we present the re-emergence of Embden-Meyerhof-Parnas glycolysis through promiscuous 1-phosphofructokinase (FruK) recruitment in a strain devoid of all glycolytic routes. We show that FruK, which canonically phosphorylates F1P to FBP, can also phosphorylate F6P, effectively replacing 6-phosphofructokinase. While FruK wildtype enzymes already have basal activity on F6P and allow growth of PZ based strains when overexpressed, we show that FruK mutants with improved kinetic parameters for the promiscuous reaction enable faster growth. Growth rates of the different FruK variants correlated best with turnover rates of the FruK variants with an R^2^ of 0.85 for the *E. coli* FruK variants and an R^2^ of 0.99 for the *C. glutamicum* variants, while *K*_M_ and catalytic efficiency were less good predictors (Supplementary Fig. 11). These results are in agreement with the logic of the PZ deletion scheme. Since G6P and F6P are expected to accumulate in the PZ strains due to the deletions of *pfkAB* and *zwf* the cytosolic F6P concentrations are probably saturating or approaching saturation for most of the FruK variants, favoring differences in turnover rate during selection.

Notably, our results show that the best variant FruK K95H does not only improve F6P activity, but also F1P activity by 26%, which raises the question why *E. coli* FruK does not natively harbor the K95H mutation. In fact, the phylogenetically related enzyme PfkB carries a histidine residue at position 95 (EcFruK numbering, Fig. 5), while *C. glutamicum* FruK1 harbors a threonine. Our structures offer a steric explanation. K95 engages in a hydrogen bond with the hydroxyl group at the C2 of the fructofuranose ring of F1P and also excludes F6P from binding by limiting the space in the active site. In order for F6P to be accommodated the fructofuranose ring needs to bind in a “flipped” orientation in comparison to F1P. Substitutions of K95 with shorter residues admit an additional water molecule into the active site, providing an anchor point for the C2 hydroxyl group of F1P and also leaving enough “wiggle room” for F6P to enter and bind. Although K95T left the turnover number for F1P essentially unchanged (180 compared to ∼200 s⁻¹ for the wildtype), it raised the apparent *K*_M_ nine-fold (1,850 compared to ∼200 µM), which makes sense as it provides even more space during substrate binding, and thus likely affects *K*_M_ more than the speed of the reaction. In contrast, the Histidine substitution appears to provide a relatively closed and rigid active site architecture in comparison to the threonine substitution, while still permitting the additional water molecule to bind. While this also leads to a decreased catalytic rate on the native substrate, this is compensated by an even greater drop in *K*_M_ for both substrates, leading to the aforementioned overall increase in catalytic efficiency (Fig. 4C,D, Tab. 3). We propose this as a case of the still quite underappreciated principle of negative catalysis^24–26^, in which an enzyme evolves to disfavor an accessible side reaction. That *E. coli* FruK retains K95, and with it forgoes the 26% improvement in native catalytic efficiency that K95H could provide, suggests that the suppression of the reaction with F6P is itself the selected trait and that its benefit lies outside catalysis.

One such benefit may be regulatory. The ligand of the global transcriptional regulator Cra has recently been shown to be F1P and not FBP^27–30^, with earlier reports of FBP attributed to contamination with F1P^28^, and FruK both generates F1P through its reverse reaction and interacts physically with Cra^30^. Residue 95 could then separate the 1- and 6-phosphofructokinase reactions in *E. coli* and, with them, glycolytic flux from transcriptional control. In *C. glutamicum*, the analogous repressor SugR also responds to F1P and controls the operon encoding FruK1^31^, despite being unrelated to Cra in sequence. *C. glutamicum* cannot use residue 95 in the same way, since FruK1 already carries a threonine there. Its activity instead appears constrained at the level of turnover, where the single substitution G104A raised *k*_cat_ on the native substrate seven-fold, from 20 to 141 s⁻¹.

The *pfk* and *zwf* deletions appear to have disrupted this regulatory balance, causing hexose-phosphate stress, a known source of toxicity in *E. coli*^17,32^. Both organisms responded with the same two-part solution. First, *fruK* was released from repression by Cra or the unrelated SugR, then the enzyme was refined through active site mutations. In *E. coli*, truncation of Cra raised FruK levels more than hundred-fold (Fig. 3D)^20^ before substitutions at position 95 were selected on glucose; in *C. glutamicum*, three of four suppressors carried *sugR* mutations alongside L29F or G104A. Consistent with this, deleting *fruK* restored the stress and was tolerated only after an additional truncation of PTS enzyme 1 (E422*), which is known to enable pan-tolerance to various stresses^33^.

Our results stand in contrast to previous attempts at measuring FruK activity on F6P^34,35^, but are in good agreement with the cross reactivity reported for 6-phosphofructokinases on F1P and with the promiscuity of ribokinase family enzymes in general^36–38^. PfkA of *E. coli* accepts F1P, albeit with 60-fold lower activity than its native substrate^38^, and FruK of the halophilic archaeon *Haloferax volcanii* accepts F6P with 36-fold lower activity^39^. While PfkB has not yet been shown to accept F1P, its promiscuity to tagatose-6-phosphate^36^ and similarity to our FruK-K95H mutant active site would suggest it could catalyze that reaction.

The substitutions recovered in our selections are moreover part of extant sequence diversity. Our phylogeny resolves FruK into subclades that differ at position 95, with both histidine and threonine occurring naturally (Fig. 5). FruK-K95T discriminates F1P over F6P ∼61-fold against ∼65-fold for *C. glutamicum* FruK1, while FruK1 retains roughly 10-fold higher catalytic efficiency on both. A single substitution in *E. coli* thus seems to have reproduced the specificity of the natural threonine subclade. Our recovered second-site substitutions, seem to modulate the effect of the threonine substitution, since G110S and T34I both roughly double the turnover number for F6P (6.8 to 12.3 and 15.7 s⁻¹) while halving the apparent *K*_M_ (4,180 to 1,660 and 2,100 µM), together raising catalytic efficiency on the new substrate about 4.6-fold. This narrows the difference in catalytic efficiency to *C. glutamicum* FruK1 from roughly 10-fold to about two-fold, showing how changes in parts of the enzyme that are further away from the active site approach the performance of the natural threonine scaffold. That *C. glutamicum* and *E. coli* converged on mutations in the same enzyme under an equivalent constraint suggests that the FruK bypass is broadly accessible across the Gram divide.

Strains comparable to the ones presented here have recently been used to engineer autotrophic growth via the CBB cycle^40–42^. In those works, activity of the phosphoribulokinase and RuBisCO bypass allowed for immediate growth of the strain, which may not have left the FruK bypass sufficient time to evolve. More generally, our results suggest a criterion that may be transferable to other systems. An active site residue that is maintained against the catalytic interest of its own enzyme marks a position at which a latent reaction is actively suppressed, and such positions may be a useful place to look for accessible new functions in other enzyme families. Promiscuous enzymes illustrate the capacity of metabolism to adapt^5,6^, to acquire missing functions and even to become bifunctional, and that this remains possible between enzymes that have diverged to about 30 % sequence identity underscores the preserved plasticity of the ribokinase scaffold. Whether the latent 6-phosphofructokinase activity of a dedicated fructose kinase also marks the route by which EMP glycolysis was originally assembled from underground metabolism remains speculative, but our results show that the transition requires remarkably little.

## Materials and Methods

### Strains and genomic modifications

All strains employed in this study are listed in Supplementary table 2. *E. coli* SIJ488 (a lambda red system expressing derivative of E. coli MG1655) served as the base strain for all genomic modifications and as wildtype reference wherever indicated. Self-made DH5a lambda pir cells were used for cloning. *E. coli* BL21 (DE3) was used for protein production. Genetic knockouts in *E. coli* were generated with lambda red engineering and P1 phage transduction. Knockout cassettes for lambda red engineering were generated from pKD3 (Chloramphenicol resistance) and pKD4 (Kanamycin resistance) plasmids while attaching 50bp homology arms targeting the gene of interest. 400-800 ng of KO-cassette DNA was electroporated into washed and cooled SIJ488 cells that were induced 45 min prior with 15mM L-arabinose at OD=0.3. Positive gene knockouts were confirmed by PCR. Chloramphenicol and Kanamycin resistance cassette were recycled via the flippase reaction. Cells were induced cells at OD=0.2 with 50mM L-Rhamnose, cultivated in LB media without antibiotics at 30°C and plated on LB plates without antibiotics. P1 phage lysates were prepared using strains from the Keio collection or lambda red derived knockout strains as donors. In brief, 3mL of mid-log grown donor cells in LB + 5mM CaCl2 were lysed with 80µL starter lysate over the course of 5 hours, and the lysate filtered through a 0.2 µm filter. Transduction was performed by growing the recipient culture to mid log, resuspending it in LB supplemented with 10mM CaCl2 and 5mM MgSO4, adding 100 µL freshly prepared donor lysate, incubating for 30min and stopping the infection process via addition of LB + citrate (65 mM final concentration). Cells were recovered for 3 hours at 37°C, washed twice in LB+100mM citrate and plated on LB media plates containing antibiotics and 5mM citrate. Colonies were picked on fresh plates with the same supplements 2 times before processing them further and saving them as glycerol stocks.

C. glutamicum WT ATCC13032 generously gifted from University of Bielefeld, was used for the generation of *pfk* and *zwf* KO strains, via the pK19mobsacB system^43^. The pJYS3 plasmid system was used for fruk1 deletion^44^. *Δpfk Δzwf Δfruk1* was generated to confirm that the strain’s growth is abolished on glucose. The suicide vector pK19mobsacB KO strategy has been described at length elsewhere^45^. The CRISPR-Cpf1-based plasmid pJYS3 encodes FnCpf1 from *Francisella novicida*, a target -specific crRNA cassette, kanamycin resistant gene, temperature-sensitive replication machinery, and variable flanking homology arms (HR1/HR2) that guide incorporation. The Cpf1-crRNA complex introduces a targeted double-strand break and facilitates homologous recombination with the plasmid-borne repair template. Following selection on LB agar containing 15 or 50 µg/µl kanamycin, plasmid curing was performed by overnight cultivation in antibiotic-free medium at 34 °C^44^. Plasmid loss was confirmed by comparing growth on kanamycin-containing and antibiotic-free LB agar. All gene deletions were subsequently verified by colony PCR.

### Cloning

The pZ-ASS-EcFruK plasmid was created in the following way. The donor pZ-ASS plasmid (pZ-ASS-Mcherry-Cap) was digested with NsiI (Mph1103I)+NheI+DpnI and treated with alkaline phosphatase to generate the backbone pZ-ASS fragment. The insert was created by PCR amplification of the fruK gene from E. coli K12 MG1655 with primers 4027+4028 followed by restriction digestion with NsiI (Mph1103I)+NheI. Insert and backbone were then ligated to form the final plasmid and transformed into *E. coli* DH5a.

For his-tag purification FruK variants were cloned into pET16b vectors. To this end, FruK variants were amplified via PCR using fruK-fwd-NdeI and fruK-rev-BamHI (CGCATATGatgagcagacgtgttgctact and CGGGATCCtcagttaaaaggttgtaagtcgacgcg) while attaching restriction sites. Amplified gene inserts and miniprepped pET16b were digested using NdeI and BamHI and ligated to create the pET16b-fruK plasmids. Plasmids were sequenced using Sanger sequencing for confirmation.

### Cultivation conditions

Routine cloning and lab cultivation of DH5a and SIJ488 strains was conducted in LB or on LB agar media (5 g/L yeast extract, 10 g/L tryptone, 10 g/L NaCl and 15 g/L agar if applicable). Growth experiments were performed in M9 minimal media (50mM Na_2_HPO_4_, 20mM KH_2_PO_4_, 20mM NH_4_Cl, 2mM MgSO_4_, 1mM NaCl, 134 μM EDTA, 100μM CaCl_2_, 13μM FeCl_3_·6H_2_O, 6.2 μM ZnCl_2_, 1.62 μM H_3_BO_3_, 0.76μM CuCl_2_·2H2O, 0.42 μM CoCl_2_·2 H_2_O, 0.081 μM MnCl_2_·4 H_2_O) in tubes or in 96-well plates. Carbon sources were added as indicated. Media in glass tubes was always supplemented with antibiotics, while it was omitted for experiments in the plate readers to maximize strain performance. For the latter 2 mL of cells were washed 3 times in M9 media without carbon source to remove residual carbon sources. Washed cells were diluted to a final OD of 0.01 in the 96-well microtiter plates (Nunclon Delta Surface, Thermo Scientific) in 150 μL media, topped with 50 μL mineral oil (Sigma-Aldrich) to avoid evaporation, while allowing sufficient gas exchange. Cell densities were measured at 600 nm every 10 min with intermittent orbital and linear shaking in a BioTek Epoch 2 plate reader (BioTek, Bad Friedrichshall, Germany). Using a Matlab script blank values were subtracted and OD values were converted to cuvette OD equivalents by multiplication with a factor of 4.35. Finally, a linear regression was fitted over the course of the growth curve to determine the fastest growth rate (minimal doubling time).

### Multiplex automated genome engineering (MAGE)

The Δ*pfk* Δ*zwf* Δ*fsaA* Δ*fsaB* (PFZ) strains were transformed with the pORTMAGE-2 plasmid to facilitate MAGE^46^. To reverse engineer the amino acid substitutions K95H, K95T, K95N and L265F in the *fruK* gene, oligonucleotides of 90 bps were designed using Genious 8 (Biomatters, New Zealand) and synthesized by Eurofins genomic (Luxembourg). MAGE was conducted according to Gallagher et al., 2014^47^ but cells only underwent one MAGE cycle and were then placed under selective pressure by plating on minimal plates containing 10mM glucose, minimizing screening work (Supplementary Fig. 4). The *fruK* locus was amplified from raised colonies via PCR and analyzed by Sanger sequencing (LGC Genomics, Berlin, DE). Once clones were confirmed positive for the desired FruK substitution, cells were passaged 5 times into fresh LB media without antibiotics at 30°C to cure the cells of the pORTMAGE-2 plasmid. Afterwards cells were diluted sequentially by 6 orders of magnitude and approximately 1000 cells were plated (given 1 ml of OD600=1 corresponds to 10^9^ cells) onto LB plates and incubated at 30°C. Colonies were then replica-plated onto plates containing Ampicillin and plates containing none. Positive clones were inoculated in fresh LB and cultivated at 37°C and then stored in 25% glycerol at -80°C.

### Whole genome sequencing

Genomic DNA was extracted from LB grown cells using the NucleoSpin Microbial DNA kit (MACHERY-NAGEL, Düren, Germany) according to the manufacturer’s instructions. DNA libraries were prepared (Nextera, Illumina) and sequenced on a paired-end Ilumina NovaSeq 6000 platform (Novogene). Raw reads were trimmed and mapped to *E. coli* SIJ488 (GenBank: CP132594) or *C. glutamicum* (GenBank: NC_006958.1) as reference genome using breseq (v0.36.1)^48^.

### Quantitative reverse transcriptase-PCR (qRT-PCR)

To quantify mRNA levels cells were harvested in exponential phase (OD600 = ∼0.5) growing on M9 minimal medium supplied with either 20mM glycerol or 10mM glucose. Total RNA was extracted using the RNeasy Mini Kit (Qiagen) according to the manufacturer’s instructions. In brief 1 ml of cells at OD600=0.5 were mixed with 2 volumes of RNAprotect Bacteria Reagent (Qiagen) and lysed enzymatically. DNA was removed on-column via RNase-free DNase (Qiagen) and spin-column-based purification of RNA. RNA concentration and integrity were determined by NanoDrop and gel electrophoresis. cDNA was synthesized from 0.5 μg RNA via reverse transcriptase using the qScript cDNA Synthesis Kit (QuantaBio). Primers were designed using Quantprime (amplicon size 61 bps, Rank score 1.4567)^49^. Quantitative real-time PCR (qRT-PCR) was performed in technical triplicates using the Maxima SYBR Green/ROX qPCR Master Mix (Thermo Scientific). Melt-curve analysis was performed to exclude non-specific amplification products. The *rrsA* gene encoding 16S rRNA was chosen as a for expression normalization, since it is a well-established reference transcript^50^. Reported data represent 2-ΔΔCT values that were calculated individually per sample relative to the average of the WT replicate ΔCt(Ct(GOI)-Ct(rrsA)) values. Controls without reverse transcriptase showed that the contribution of the residual gDNA amounted to less than 3% of the signal in the PZ derived strains, while they were indistinguishable from +RT samples for the wildtype samples. Data shown were not corrected for this gDNA contamination, but gDNA signal is listed in the Supplementary information. Primer efficiencies for fruK and rrsA amplification were assessed by qPCR on a *E. coli* gDNA tenfold dilution series spanning 6 orders of magnitude (Supplementary Fig. 5). Samples were run in triplicates. A linear regression was fitted to the average of the data to generate the slope. Primer efficiency was calculated as E = (10^(−1/slope)^ - 1) * 100 and showed that the fruK and rrsA primers had 99.9 and 98.4% efficiency respectively.

### Phylogenetic analysis

Trees were built through iterative blastp searches starting from the *E. coli* FruK sequence, selection of diverse sequences based on % identity to the *E. coli* sequence, sequence alignment, phylogeny generation, and re-blasting with sequences in interesting positions or with different residues at the active site (identified via position in sequence alignment). Additionally, sequences from known protein families related to FruK were used to blastp & Uniprot search for more representatives to then include those in the phylogeny. Protein families were sourced from Zhang et al., 2004^51^ and Cabrera et al., 2010^21^. Sequences were aligned using MUSCLE v5 and trees were built using RaxML-ng and the LG+G substitution model (command: raxml-ng --msa ribokinase_family_v1.afas --model LG+G --prefix T1 --threads 2 -- seed 2 –redo). For fast iteration phylogeny building (during sequence selection) “FastTree” was used. Active site positions of interest were determined to be positions 15, 41, 91, 93, 95, 110, 185, 255 (*E. coli* numbering). Sequence logos were created using Weblogo 3.7.12 after reducing multiple sequence alignments of the sequences in each clade to only the active site positions. For example: for the PfkB clade, all PfkB sequences were put into a separate sequence alignment and all columns of residues except for those at position 15, 41,… etc, were deleted. Highlighted sequences are roughly indicated with an asterisk (*) in the phylogeny. Figures were assembled in Adobe Illustrator. PdxK was included as an outgroup as it was previously described to be more ancient than FruK^21^.

### Enzyme purification

*E. coli* BL21 (DE3) cells (1 Liter cultures) containing the expression vectors encoding His_10_-tagged versions of the different FruK variants were grown overnight at 25°C in TB medium with 100 µg/L ampicillin and without applying any induction, relying solely on leaky expression by the T7 promotor. Cells were harvested at 8°C and 7,500x g. The cell pellets were stored at -70 °C before lysis. Cells were resuspended in a three-fold volume of lysis buffer (1 g of cells in 3 mL of buffer) containing 50 mM MOPS/KOH (pH 7.4), 300 mM KCl, and 1 M L-proline. Proline was added to avoid precipitation of the enzyme, which was observed in buffers lacking proline. Before lysis 0.1 mg mL-1 DNase was added to the resuspended cells. The cells were then passed 3 times through an LM10 microfluidizer (Microfluidics, USA) with interaction chamber H10Z at 18,000 psi. Cell debris was removed by centrifugation at 100,000 × *g* and 8 °C for 1 h. The supernatant was applied at a flow rate of 1 mL min_-1_ to a 1 mL HisTrap™ FF column (Cytiva, Germany), which had been equilibrated with 5 column volumes of lysis buffer. The column was washed with 20 column volumes of lysis buffer containing an additional 100 mM imidazole. The protein was eluted with lysis buffer containing 500 mM imidazole. Fractions containing the eluted enzyme were pooled and concentrated, before they were applied to a HiLoad 16/600 Superdex 200 pg size exclusion column (Cytiva, Germany) at a flow rate of 1 mL min_-1_. This column had been equilibrated with buffer containing 20 mM MOPS/KOH (pH 7.4), 75 mM KCl, and 1 M L-proline. FruK variants eluted as tetramers from the size exclusion column. The protein was then again concentrated and either directly used for crystallization or stored at -20 °C in the presence of 20 % glycerol for subsequent enzyme activity assays.

### Enzyme assays

A spectrophotometric assay was used to measure enzyme activities at 37 °C in 1 cm quartz cuvettes. The assay mix (100 µL) comprised 200 mM MOPS/KOH (pH 7.8), 2 mM ATP, 4 mM phosphoenolpyruvate, 5 mM MgCl_2_, 0.5 mM NADH, as well as pyruvate kinase, and lactate dehydrogenase as coupling enzymes. The assay mix was complemented with varying amounts of fructose-1-phosphate or fructose-6-phosphate, as well as appropriate amounts of FruK variants to determine catalytic activities and apparent *K*_M_-values. ADP formed by the kinase activity was re-converted to ATP by pyruvate kinase, consuming phosphoenolpyruvate and producing pyruvate in the process. Pyruvate was then converted into lactate by lactate dehydrogenase, consuming NADH, the latter was monitored at 365 nm. A molar extinction coefficient ε_365_ for NADH of 3330 M^-1^ cm^-1^ was used to calculate enzyme activities.

To better understand the apparent mutase activity during FruK crystallization we applied an LC-MS based assay. FruK wild type enzyme as well as FruK variant K95H (1.5 µg, respectively) were incubated for 30 minutes in 50 mM MOPS/KOH buffer (pH 7.8), 5 mM MgCl_2_, 2 mM of either ATP or ADP, and 1 mM of either F1P, F6P, or FBP. The total reaction volumes were 0.2 mL. The reactions were started by addition of enzyme and samples of 20 µL were withdrawn after 30 minutes. A reaction mixture without enzyme served as a negative control. Samples were immediately stopped by mixing with 60 µL of cold 6.5 % (v/v) formic acid. Samples were spun down for 15 minutes at 4°C and 17,000x g to remove precipitated protein. The supernatants were diluted 1:10 in water and applied to LC-MS analysis. Determination of F1P, F6P, and FBP was performed using a HRES-LC-MS. The chromatography was performed using a Thermo Scientific Dionex Integrion HPIC system equipped with an EGC 500 KOH eluent generator, and a Dionex ADRS 600 suppressor operated in constant-current mode at 95 mA. Separation was achieved on a Dionex IonPac AS11-HC-4µm column (150 × 2 mm) equipped with a corresponding guard column (50 × 2 mm), using an eluent flow rate of 0.38 mL/min and a regenerator flow rate of 0.75 mL/min (water). The column temperature was maintained at 30 °C. The injection volume was 5 µL, using an automated sampler. The programmed KOH gradient was as follows: 0–7.5 min, 15– 32 mM; 7.5–7.6 min, 32–100 mM; 7.6–14 min, 100 mM; 14–14.1 min, 100–15 mM; and 14.1– 15 min, 15 mM. A Thermo Scientific ID-X Orbitrap mass spectrometer was used in negative mode with an electrospray ionization source and the following conditions: ESI spray voltage 3300 V, sheath gas at 60 arbitrary units, auxiliary gas at 15 arbitrary units, sweep gas at 2 arbitrary units, ion transfer tube temperature at 300°C and vaporizer temperature at 300 °C. Compounds were detected in a targeted mode and identified based on their exact mass with a mass window of 5 ppm and retention time compared to standards. Extracted ion chromatograms of the [M-H]-forms were integrated using Tracefinder software (Thermo Scientific). Concentrations were calculated based on an external calibration curve.

### Crystallization

The sitting-drop vapor-diffusion method was used for all crystallization attempts. Crystals were grown at 16 °C.

For crystallization of the wild type variant of FruK, the enzyme (3.4 mg/mL) in 20 mM MOPS/KOH buffer (pH 7.4) containing 75 mM KCl and 1 M L-proline was mixed in a 1:1 ratio with crystallization condition comprised of 1 M (NH_4_)_2_SO_4_, 75 mM TRIS/HCl (pH 8.5), 150 mM Li_2_SO_4_, 2 mM fructose-1-phosphate, 4 mM ADP, and 10 mM MgCl_2_. The final drop volume was 0.8 microliters. Prior to flash freezing the crystals in liquid nitrogen, the mother liquor was supplemented with 35 % (v/v) glycerol.

The FruK variant K95H (3.4 mg/mL) in 20 mM MOPS/KOH buffer (pH 7.4) containing 75 mM KCl and 1 M L-proline was mixed in a 1:1 ratio with crystallization condition comprised of 1 M (NH_4_)_2_SO_4_, 75 mM TRIS/HCl (pH 8.5), 150 mM Li_2_SO_4_, 8 mM fructose-6-phosphate, 4 mM ADP, and 10 mM MgCl_2_. The final drop volume was 0.8 microliters. Prior to flash freezing the crystals in liquid nitrogen, the mother liquor was supplemented with 30 % (v/v) glycerol. Although crystals were grown in the presence of fructose-6-phosphate we could only observe electron density accommodating fructose-1-phosphate in the active site (see Results section). An additional soaking step in 8mM fructose-6-phosphate of about 1 minute was applied before freezing the crystals in order to exchange the bound substrate. The mother liquor was complemented with 25 mM fructose-6-phosphate and 12 mM MgCl_2_ before glycerol was added to a final concentration of 30 % (v/v).

The FruK variant K95T/G110S (3.5 mg/mL) in 20 mM MOPS/KOH buffer (pH 7.4) containing 75 mM KCl and 1 M L-proline was mixed in a 1:1 ratio with crystallization condition comprised of 1 M ammonium sulfate, 75 mM TRIS/HCl (pH 8.5), 150 mM lithium sulfate, 8 mM fructose-6-phosphate, 4 mM ADP, and 10 mM MgCl_2_. The final drop volume was 0.8 microliters. Prior to flash freezing the crystals in liquid nitrogen, the mother liquor was supplemented with 30 % (v/v) glycerol. Although crystals were grown in the presence of fructose-6-phosphate we could only observe electron density accommodating fructose-1-phosphate in the active site (see Results). An additional soaking step of about 1 minute was applied before freezing the crystals in order to exchange the bound substrate. The mother liquor was complemented with 20 mM fructose-6-phosphate, 2 mM ADP, and 5 mM MgCl_2_ before glycerol was added to a final concentration of 30 % (v/v). The soaking resulted in a change in electron density within the active site of the mutant variant, which allowed the modeling of fructose-1,6-bisphosphate, X-ray diffraction data (Supplementary table 1) were collected at beamlines PETRA III P13 and P14 of DESY (Deutsches Elektronen-Synchrotron, Hamburg) as well as at the beam lines ID23-2 and ID30B of the ESRF (European Synchrotron Radiation Facility, Grenoble). Data sets were processed with the XDS software package^52^. Structures were solved by molecular replacement using Phaser of the Phenix software package (v1.21)^53^, built with Phenix. Autobuild and refined with Phenix.Refine. Additional modelling, manual refinement, ligand fitting, and water picking was done in Coot (v.0.9.8.3)^54^. Final positional and B-factor refinements were performed using Phenix.Refine. Structural models for the FruK wild type variant, as well as the two FruK mutant variants with different bound substrates/products were deposited in the Protein Data Bank of Europe (PDBe) under PDB accessions 9QV0, 9QV1, 9QV2, 9QV3, and 9QV4, respectively (Supplementary table 1). Figures were made using PyMOL Molecular Graphics System (version 2.5.7; Schrödinger) and PoseEdit^55^

## Acknowledgments

The authors dedicate this work to Arren Bar-Even, who was a brilliant visionary in the field of synthetic biology, a precious friend to many of us and who was involved in the initial findings of this study. This study was funded by the Max Planck Society (to B.D., J.Z., A.S., L.S., A.B.E., T.J.E. and S.N.L). S.N.L, D.B., I.P., S.H., E.S. and L.T-B. acknowledge support from the BMBF funded projects ForceYield and ForceYield 2.0. The authors thank Filippo Federici for assistance in whole genome sequencing analysis.

## Author contributions

B.D. and J.Z. contributed equally. B.D., A.B.-E., T.J.E. and S.N.L. conceived the study. B.D., J.Z., and S.N.L. designed the experiments. B.D., I.P., A.S., and S.N.L. performed strain engineering, WGS analysis and growth experiments in *E. coli* and D.B., I.P., S.H., E.S., L.T.-B. experiments in *C. glutamicum*. B.D. conducted qRT-PCR analysis. J. Z. measured enzyme kinetics, performed LC/MS experiments and crystallography. L.S. performed phylogenetic analyses. A.B.-E., T.J.E. and S.N.L. supervised research. B.D., J. Z., T.J.E. and S.N.L. wrote the manuscript with contributions from all authors.

## Data availability

All data supporting the findings of this study are accessible within the paper and the accompanying Supplementary Information. Source data were provided with this paper. All strains, plasmids and materials mentioned in the manuscript can be obtained from the corresponding authors upon reasonable request.

