## Supplementary information for "Gain-of-function mutations in 1-phosphofructokinas*e (FruK)* re-establish glycolysis in bacteria"

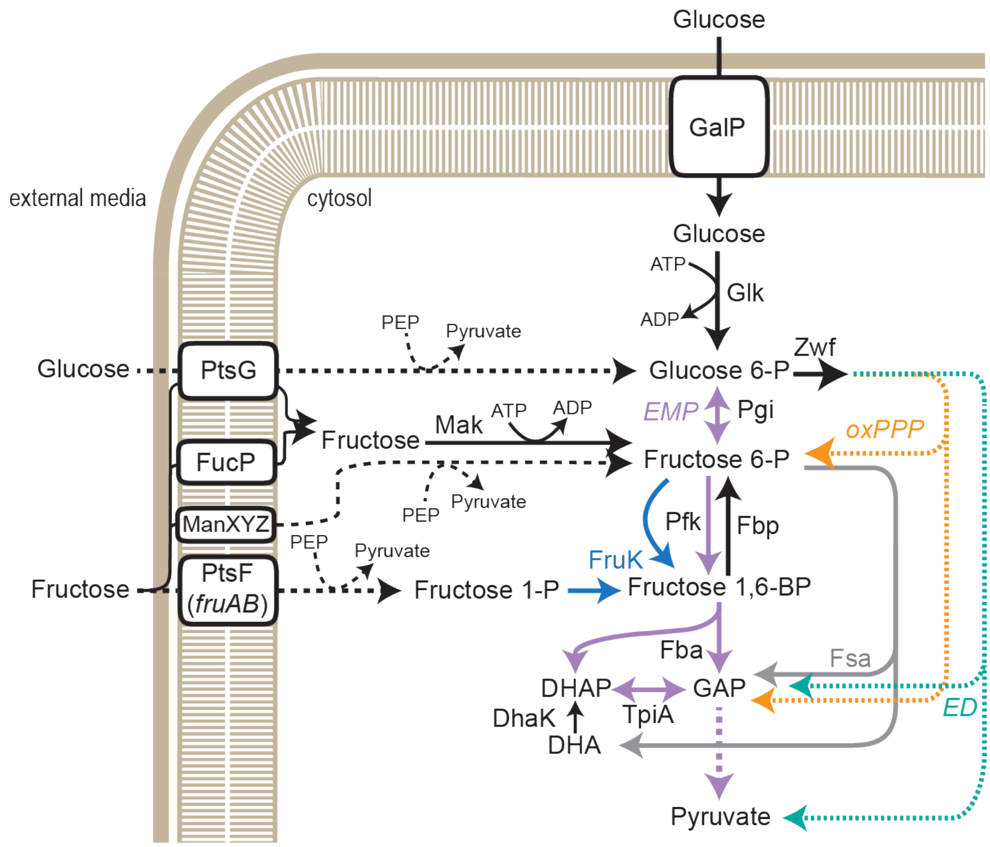


**Supplementary figure 1: Main glucose and fructose degradation routes in *E. coli* extended.** Glucose and fructose import and degradation to pyruvate are shown. Glucose and fructose both have a dedicated PTS system (PTS; PtsG and FruA, supplied with phosphoryl groups through PtsH and FruB respectively, which in turn are both phosphorylated by enzyme I (PtsI)) and alternative import and activation options (GalP/Glk). Four potential utilization routes are shown for fructose[12,13]. Main glycolytic routes are labeled and color-coded, Embden-Meyerhof-Parnas (EMP) pathway in violet, Entner-Doudoroff (ED) pathway in turquoise, oxidative pentose phosphate pathway (oxPPP) in yellow and fructose-6-phosphate aldolase (Fsa) route in grey. The canonical and novel reactions of 1-phosphofructokinase (FruK) are shown in blue. Phosphoglucose isomerase (Pgi), Fructose bisphosphate phosphatase (Fbp), 6-phosphofructokinase (Pfk), Fructose bisphosphate aldolase (Fba), Triosephosphate isomerase (TpiA), Dihydroxyacetone kinase (DhaK), Dihydroxyacetone phosphate (DHAP), Glyceraldehyde-3-phosphate (GAP).

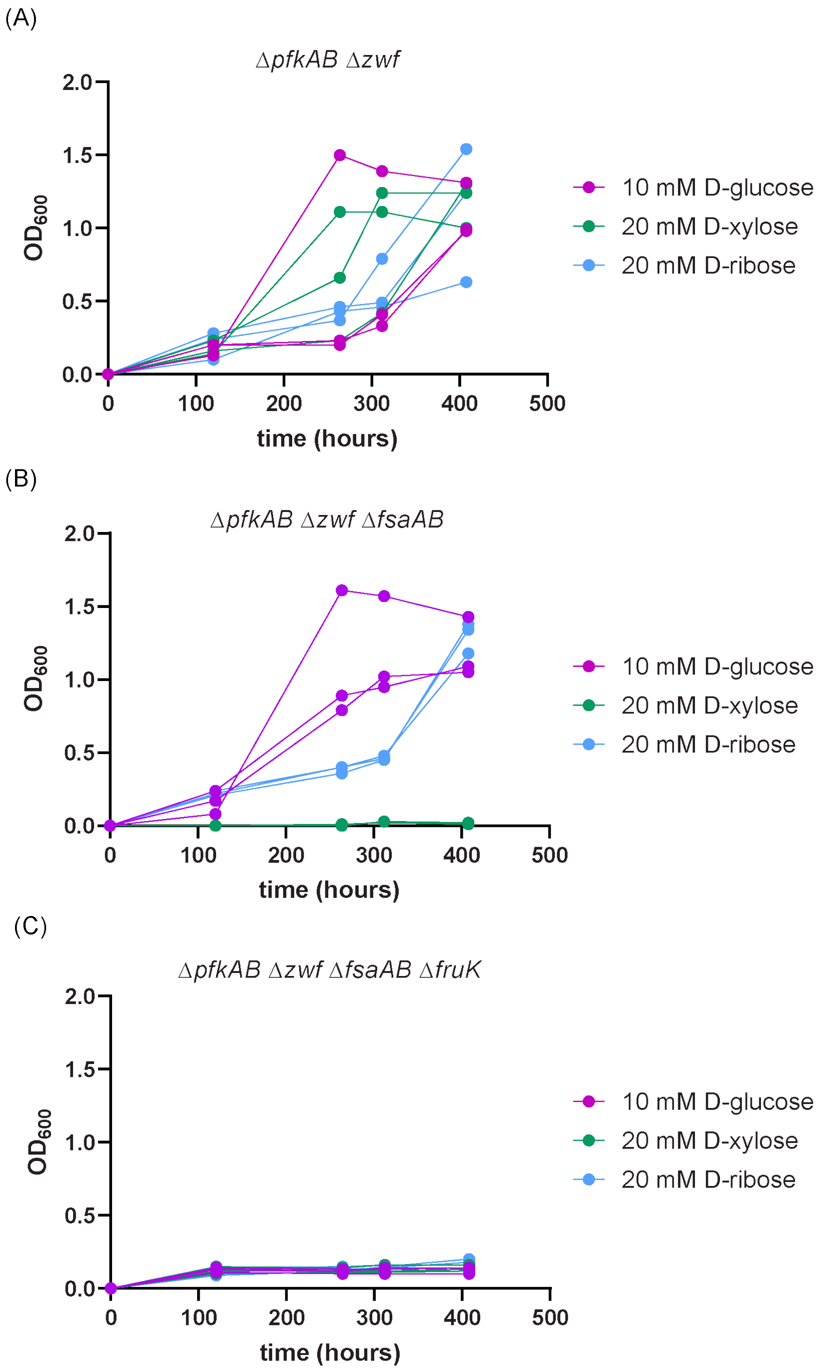


**Supplementary figure 2: Adaptive laboratory evolution experiment of glucose catabolism deficient strains of *E. coli* on different carbon sources.** Δ*pfkAB* Δ*zwf*, Δ*pfkAB* Δ*zwf* Δ*fsaAB* and Δ*pfkA* Δ*zwf* Δ*fsaAB* Δ*fruK* strains were cultivated in minimal media supplemented with 10 mM D-glucose, 20 mM D-xylose or 20 mM D-ribose over 17 days.

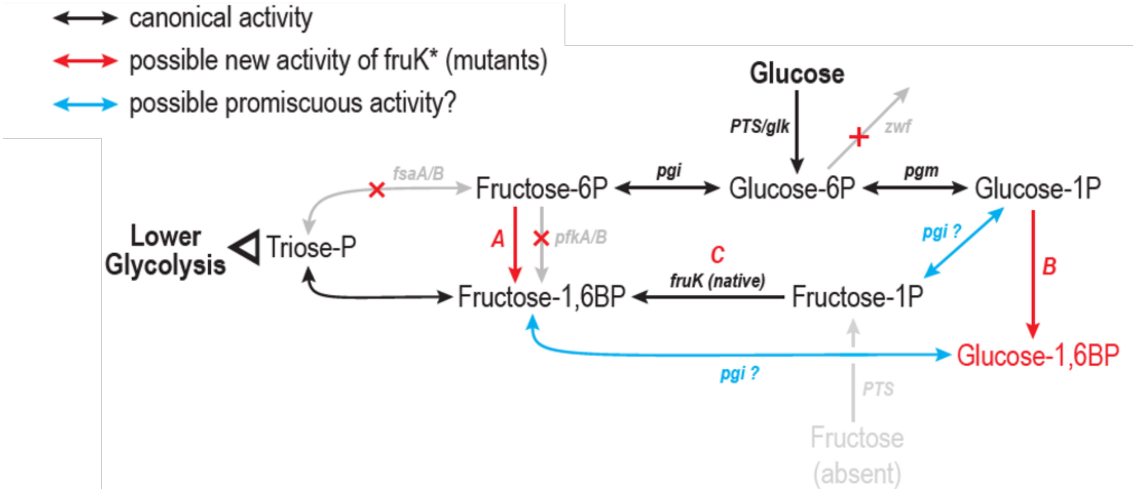


**Supplementary figure 3: Hypothetical routes for the evolved FruK bypass.** Several metabolic pathways are shown that could allow growth of the pfkAB and zwf deficient strains on glucose via mutated FruK activity in combination with wildtype phosphoglucose mutase (Pgm) and phosphoglucose isomerase (Pgi) activity.

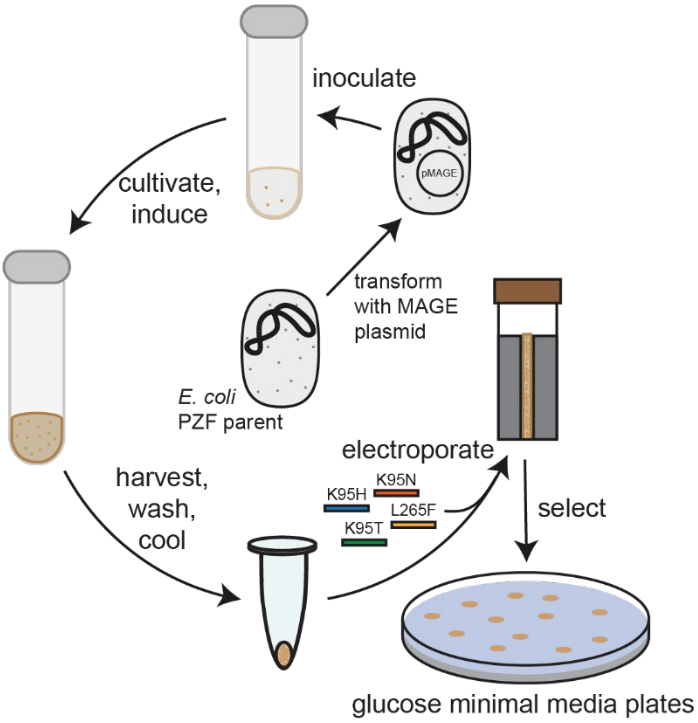


**Supplementary figure 4: MAGE-based selection workflow.** Oligos to incorporate the FruK substitutions K95H, K95T, K95N and L265F were electroporated into the parental *E. coli* PZF strain. Cells were plated on M9, 10 mM glucose agarose plates to select for cells with mutated FruK.

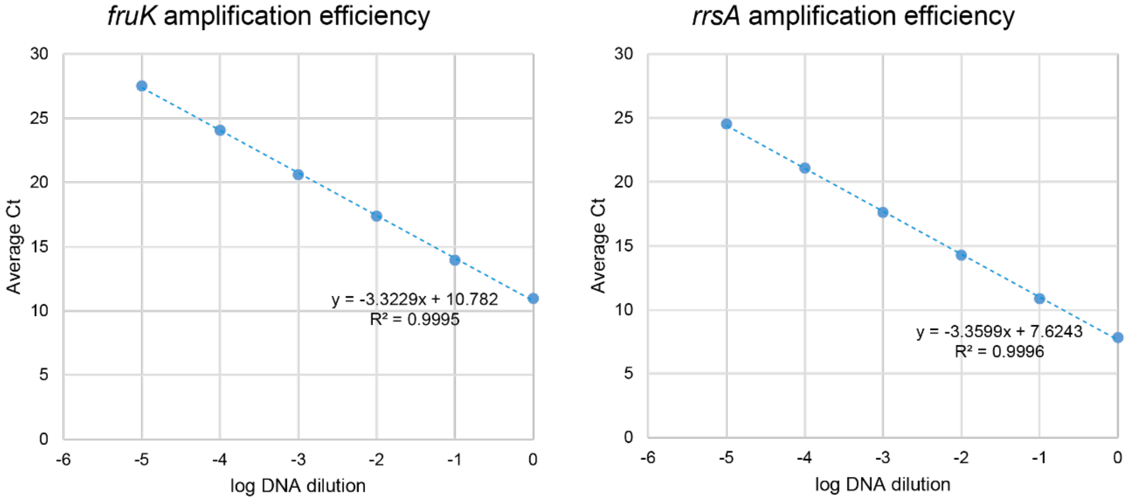


**Supplementary figure 5: Primer efficiency analysis via quantitative PCR.** Primer efficiency (E = (10(-1/slope) - 1) * 100) of the *fruK* and *rrsA* primers was determined to be 99.9 and 98.4% respectively.

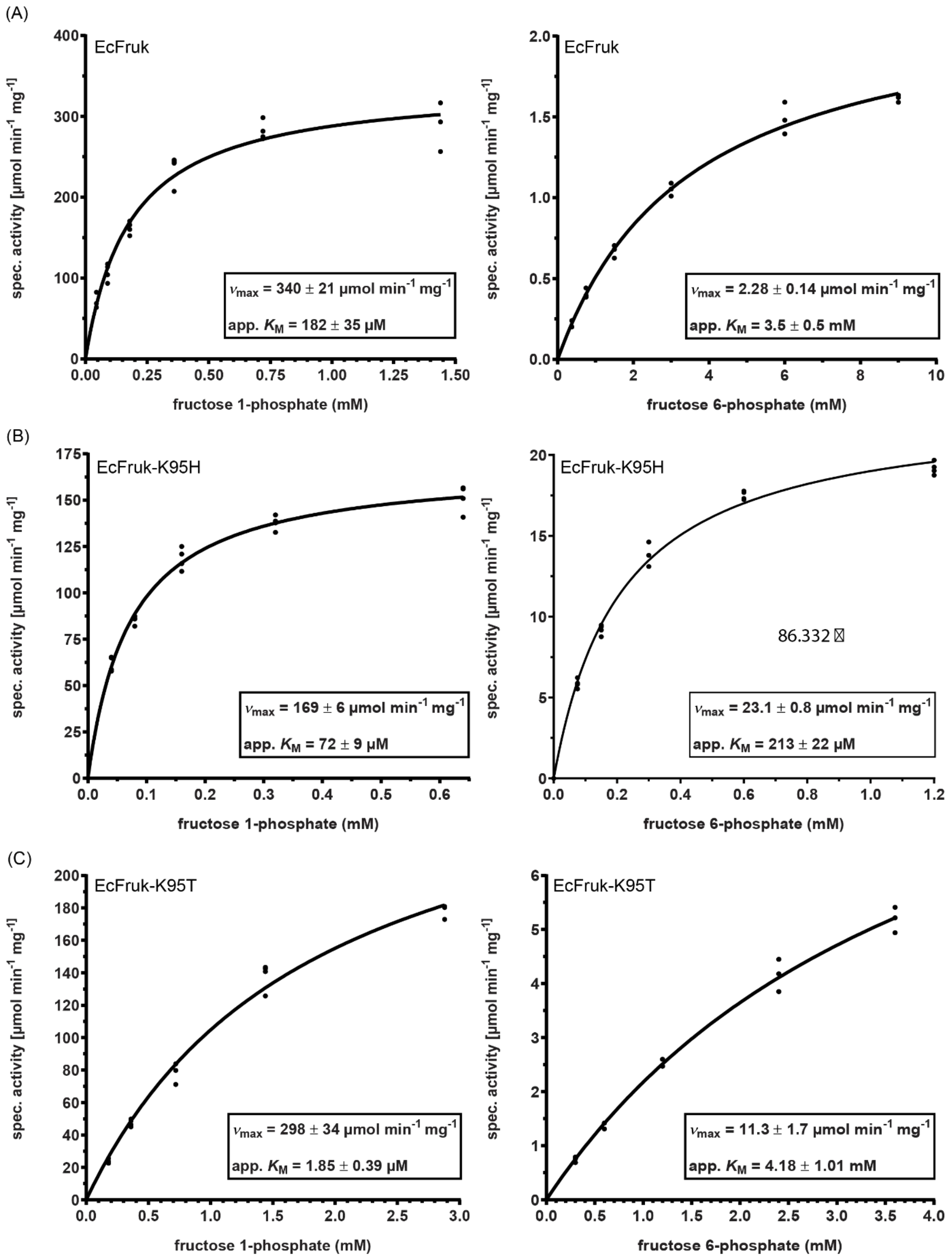
(A)
EcFruk
spec. activity [μmol min-1 mg-1]
fructose 1-phosphate (mM)
vmax = 340 ± 21 μmol min-1 mg-1
app. KM = 182 ± 35 μM
EcFruk
spec. activity [μmol min-1 mg-1]
fructose 6-phosphate (mM)
vmax = 2.28 ± 0.14 μmol min-1 mg-1
app. KM = 3.5 ± 0.5 mM
(B)
EcFruk-K95H
spec. activity [μmol min-1 mg-1]
fructose 1-phosphate (mM)
vmax = 169 ± 6 μmol min-1 mg-1
app. KM = 72 ± 9 μM
EcFruk-K95H
spec. activity [μmol min-1 mg-1]
fructose 6-phosphate (mM)
86.332
vmax = 23.1 ± 0.8 μmol min-1 mg-1
app. KM = 213 ± 22 μM
(C)
EcFruk-K95T
spec. activity [μmol min-1 mg-1]
fructose 1-phosphate (mM)
vmax = 298 ± 34 μmol min-1 mg-1
app. KM = 1.85 ± 0.39 μM
EcFruk-K95T
spec. activity [μmol min-1 mg-1]
fructose 6-phosphate (mM)
vmax = 11.3 ± 1.7 μmol min-1 mg-1
app. KM = 4.18 ± 1.01 mM

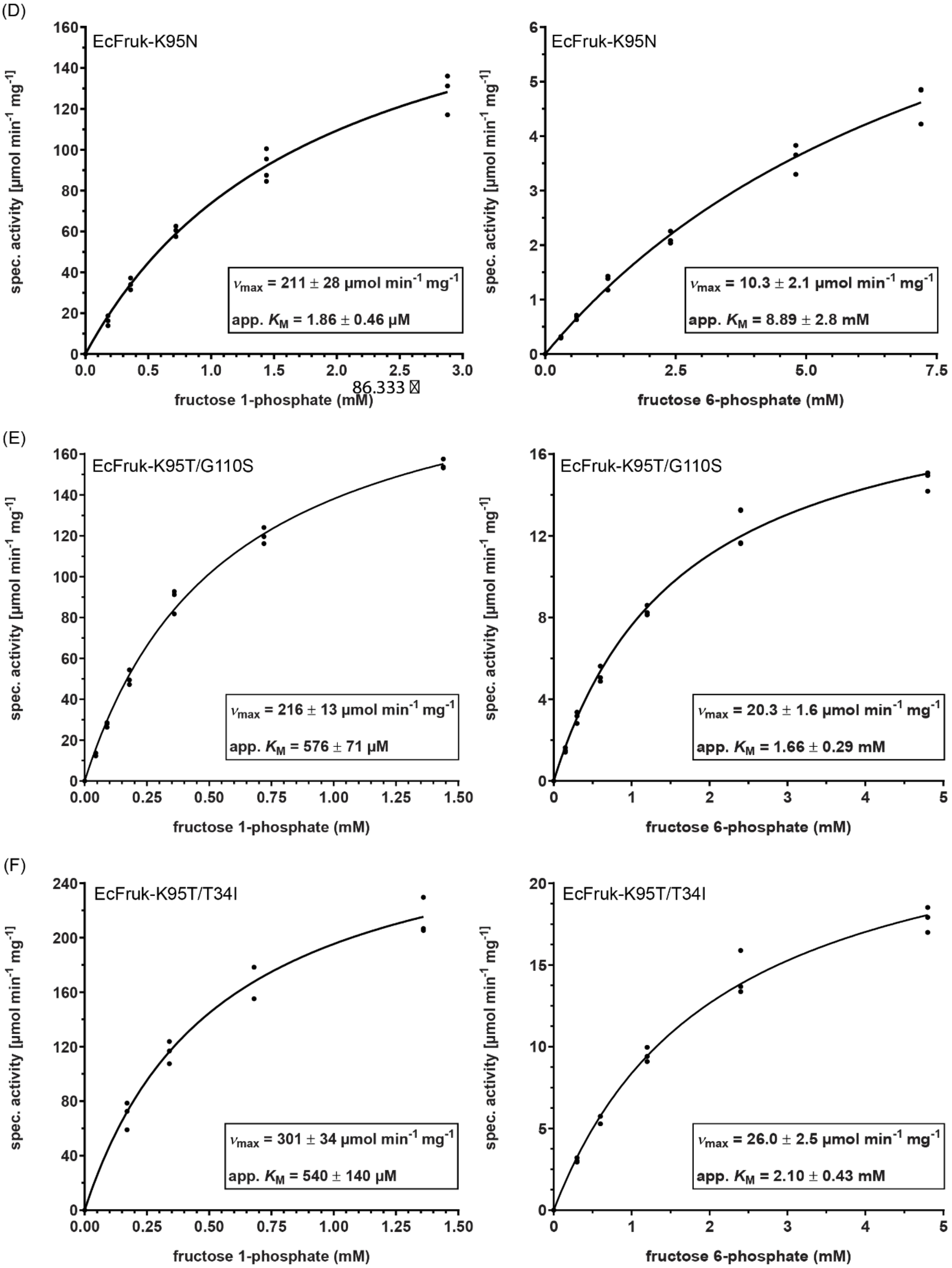

(D)
EcFruk-K95N
$v_{max}$ = 211 ± 28 µmol min$^{-1}$ mg$^{-1}$
app. $K_M$ = 1.86 ± 0.46 µM
86.333
fructose 1-phosphate (mM)
EcFruk-K95N
$v_{max}$ = 10.3 ± 2.1 µmol min$^{-1}$ mg$^{-1}$
app. $K_M$ = 8.89 ± 2.8 mM
fructose 6-phosphate (mM)
(E)
EcFruk-K95T/G110S
$v_{max}$ = 216 ± 13 µmol min$^{-1}$ mg$^{-1}$
app. $K_M$ = 576 ± 71 µM
fructose 1-phosphate (mM)
EcFruk-K95T/G110S
$v_{max}$ = 20.3 ± 1.6 µmol min$^{-1}$ mg$^{-1}$
app. $K_M$ = 1.66 ± 0.29 mM
fructose 6-phosphate (mM)
(F)
EcFruk-K95T/T34I
$v_{max}$ = 301 ± 34 µmol min$^{-1}$ mg$^{-1}$
app. $K_M$ = 540 ± 140 µM
fructose 1-phosphate (mM)
EcFruk-K95T/T34I
$v_{max}$ = 26.0 ± 2.5 µmol min$^{-1}$ mg$^{-1}$
app. $K_M$ = 2.10 ± 0.43 mM
fructose 6-phosphate (mM)
spec. activity [µmol min$^{-1}$ mg$^{-1}$]

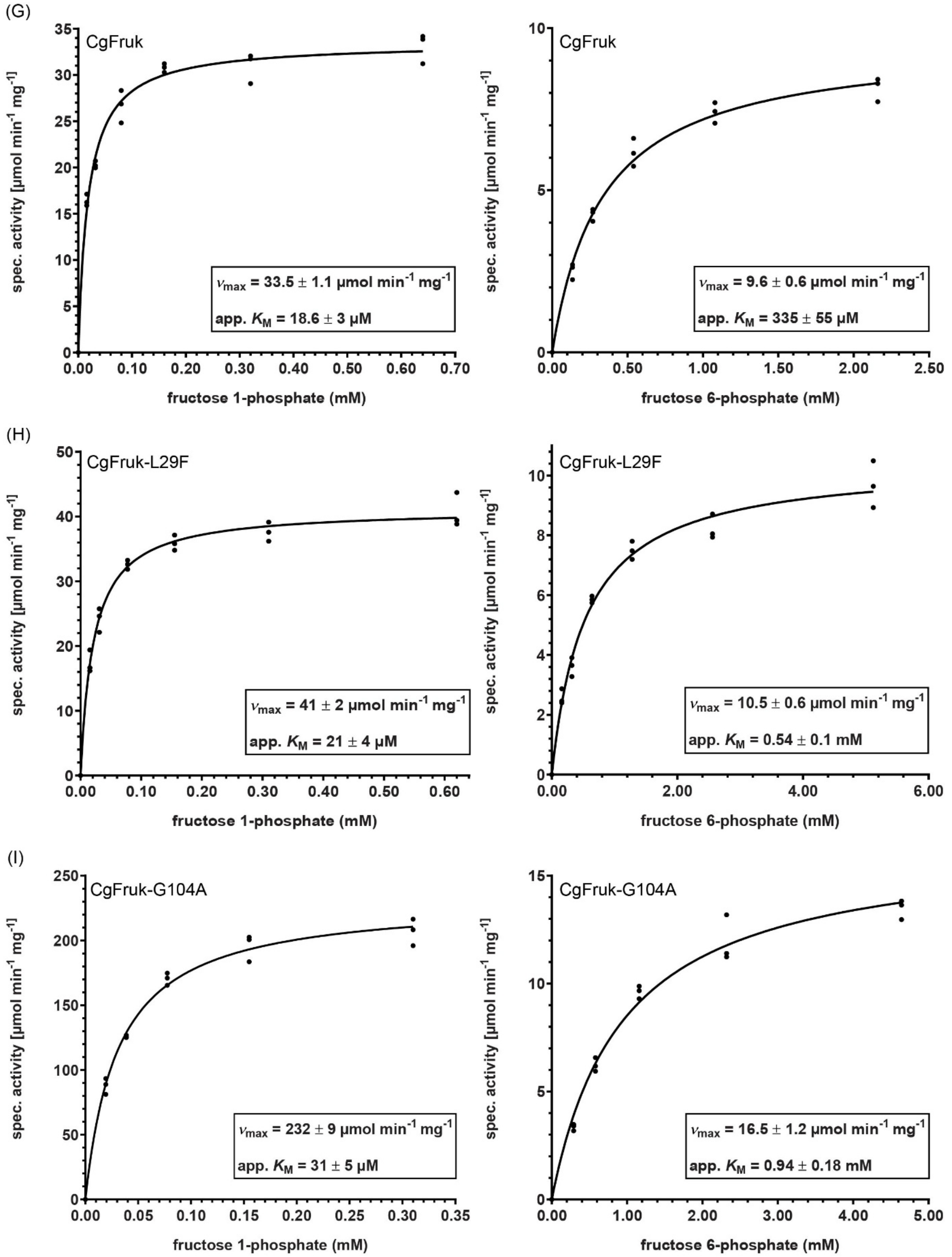


**Supplementary figure 6: Michaelis Menten kinetics of *E. coli* and *C. glutamicum* FruK variants on F1P (left) and F6P (right).** The following kinetics are shown: EcFruK wildtype (A), EcFruK-K95H (B), EcFruK-K95T (C), EcFruK-K95N (D), EcFruK-K95T/G110S (E), EcFruK-K95T/T34I (F), CgFruK1 wildtype (G), CgFruK1-L29F (H) and CgFruK1-G104A (I).

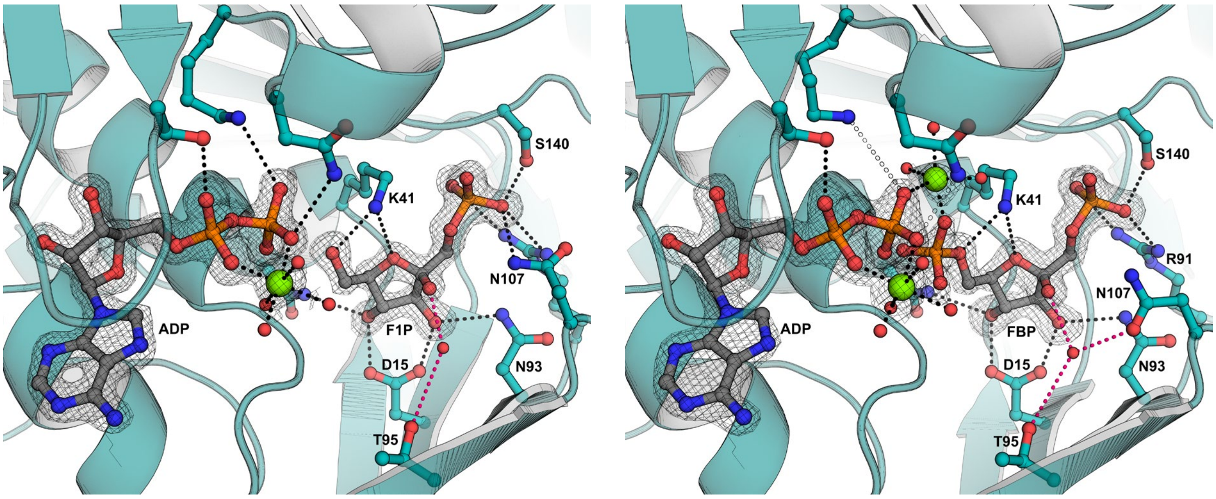


**Supplementary figure 7: Active site of the K95T/G110S variant of the *E. coli* FruK with bound substrates.** The left panel shows the active site with bound F1P, the right panel with bound FBP. Polar interactions between sidechains and ligands are depicted by dashed lines. Interactions with a water molecule occupying the space of the substituted residue K95 (here T95) are highlighted in pink. Simulated annealing omit maps are shown at a σ of 4.0 for ADP, F1P, and FBP. Note that a second magnesium ion is bound in the structure with FBP bridging the terminal phosphate group of ADP and FBP.

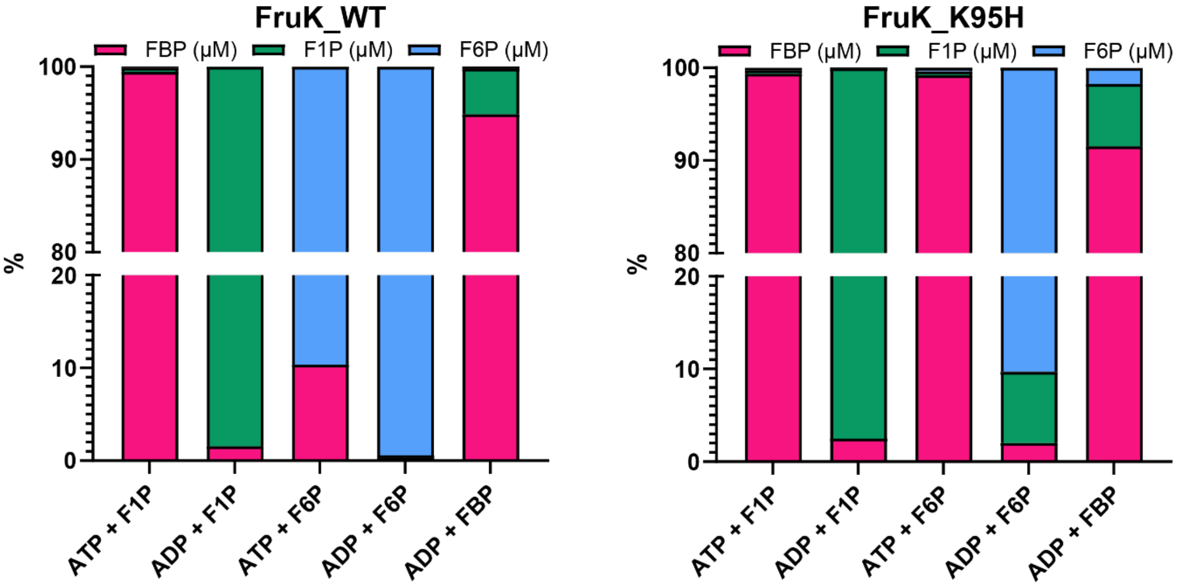


**Supplementary figure 8: Substrate and product change endpoint measurement of *E. coli* FruK variants with either F1P, F6P or FBP together with either ADP or ATP.**

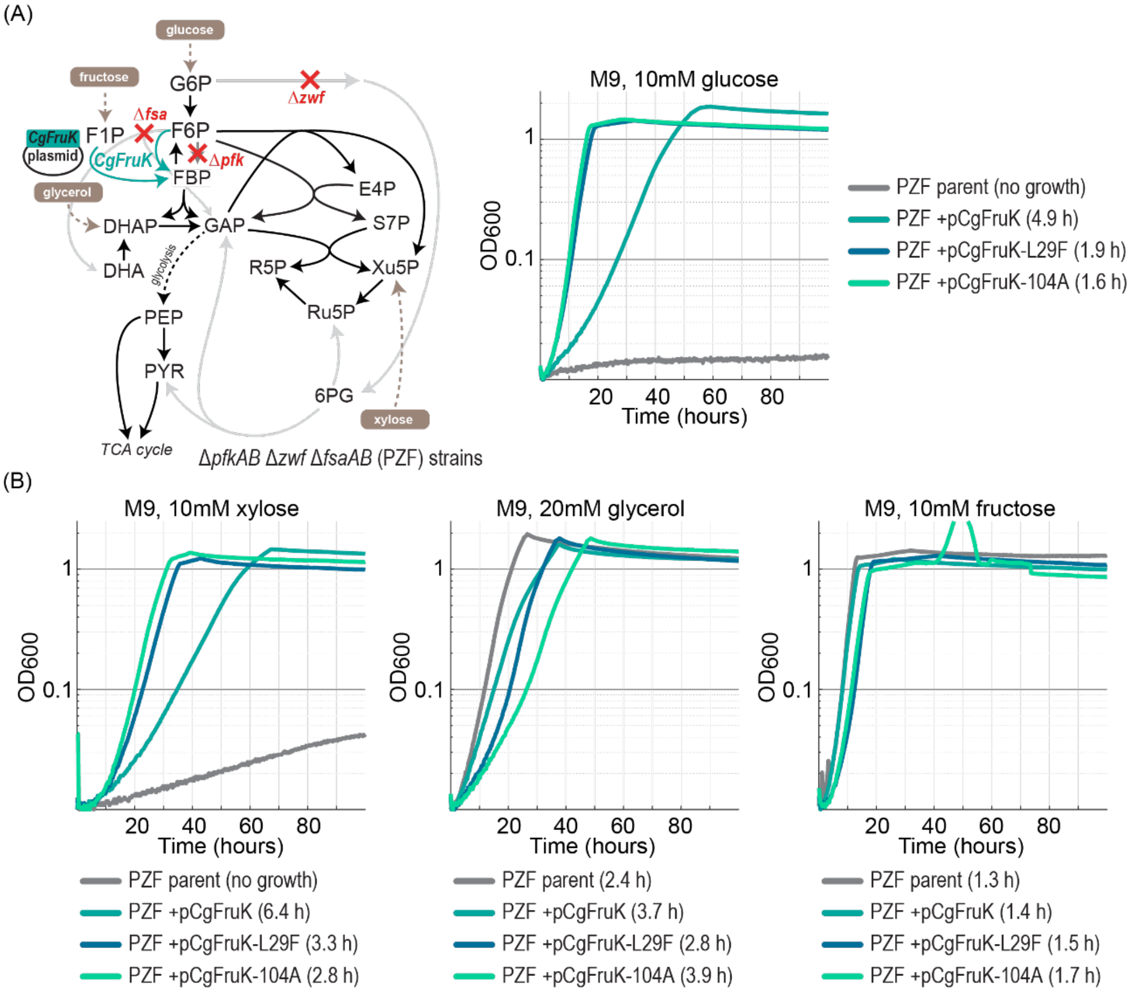


**Supplementary figure 9: Plasmid complementation experiment of the *E. coli* Δ*pfk* Δ*zwf* Δ*fsaAB* strain with *C. glutamicum* FruK1 variants restore growth on glucose and xylose.** Strains were cultivated in M9 media supplemented with either 10mM glucose, 10mM xylose, 20mM glycerol or 10mM fructose.

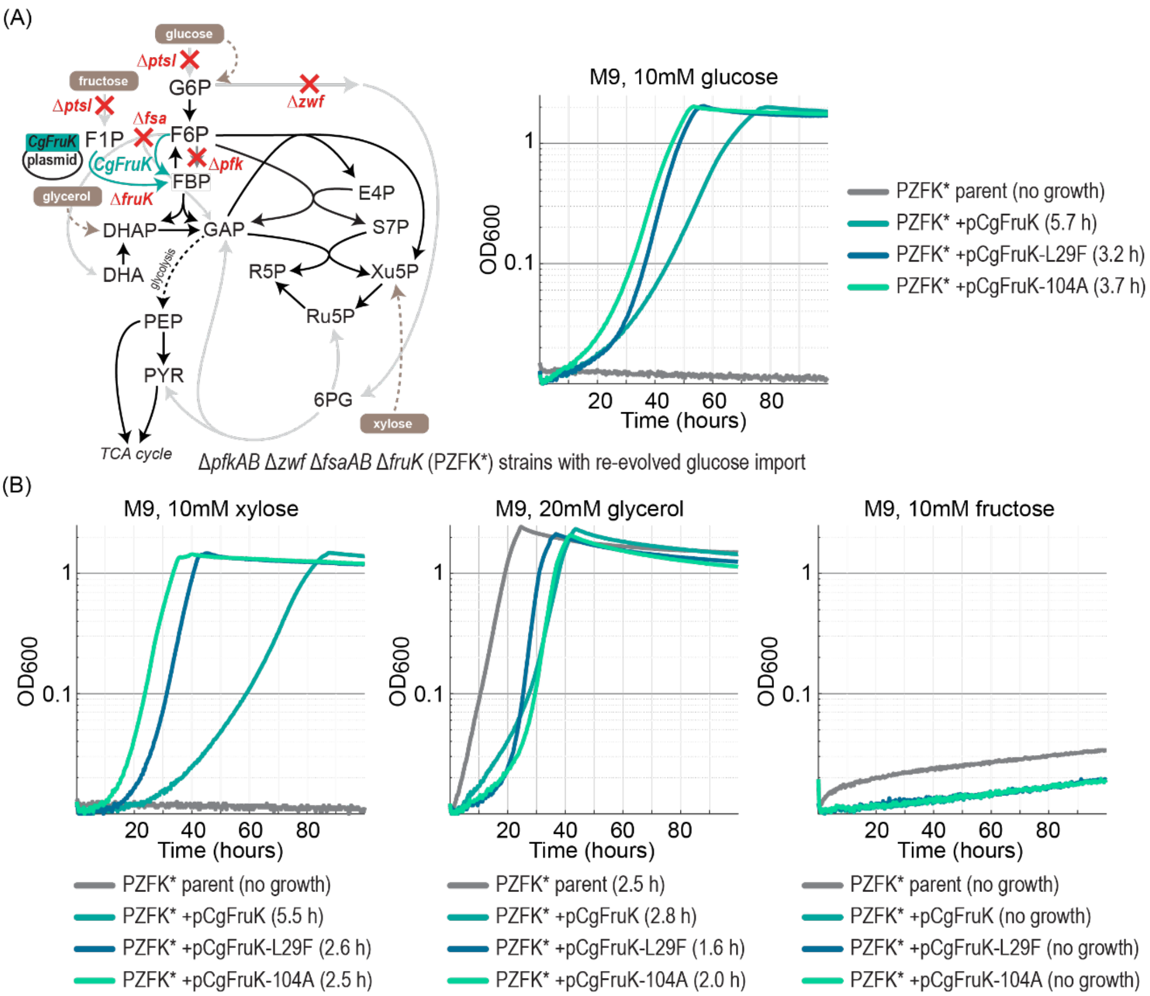


**Supplementary figure 10: Plasmid complementation experiment of the *E. coli* Δ*pfk* Δ*zwf* Δ*fsaAB* Δ*fruK* strain with *C. glutamicum* FruK1 variants restore growth on glucose and xylose.** Strains were cultivated in M9 media supplemented with either 10mM glucose, 10mM xylose, 20mM glycerol or 10mM fructose. Deletion of fruK was accompanied by truncated of ptsI (affecting canonical PTS based import of glucose and fructose. To perform the complementation test for growth on glucose we first evolved the Δ*pfk* Δ*zwf* Δ*fsaAB* Δ*fruK* strain on glucose while overexpressing FruK. This cured strain that revolved the ability to grow on glucose (potentially via GalP and Glk) was used here.

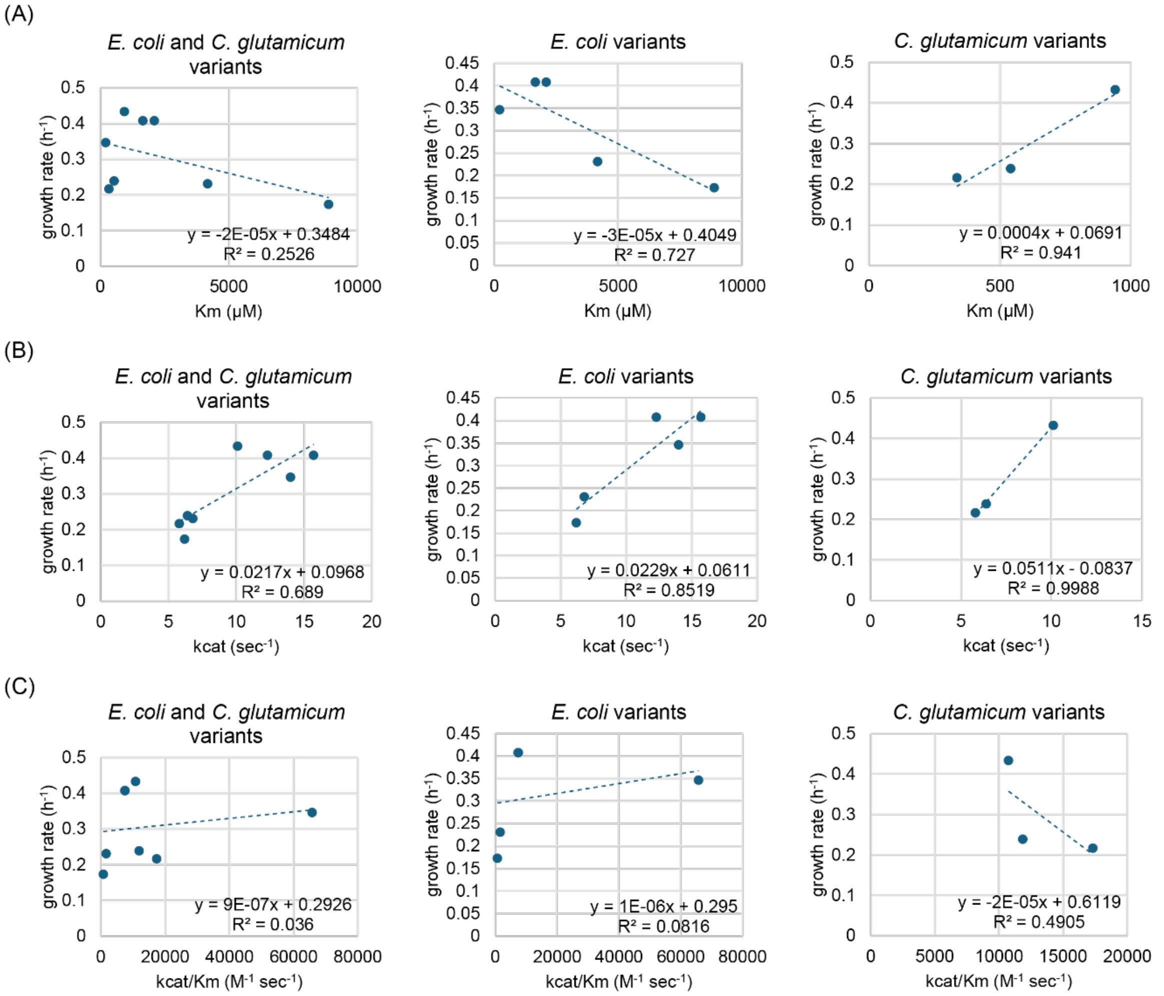


**Supplementary figure 11: Linear correlation between *E. coli* growth rates and FruK enzyme kinetic parameters.** Individual panels display the linear regression of specific growth rates of the PZF strains expressing various FruK variants on glucose plotted against the measured kinetic parameters for the apparent $K_M$ in µM, the turnover number per sec and catalytic efficiency ($k_{cat}/K_M$ in $M^{-1}$ $sec^{-1}$). Scatter points represent averages determined from triplicate measurements in 96-plate readers.

**Supplementary table 1: X-ray data collection and structure refinement statistics.**

| Accession | PDB 9QV0 | PDB 9QV1 | PDB 9QV2 | PDB 9QV3 | PDB 9QV4 |
|---|---|---|---|---|---|
| FruK variant | wild type | K95H | K95H | K95T/G110S | K95T/G110S |
| Ligands | ADP, F1X, $Mg^{2+}$ | ADP, F1X, $Mg^{2+}$ | ADP, F6P, $Mg^{2+}$ | ADP, F1X, $Mg^{2+}$ | ADP, FBP, $Mg^{2+}$ |
| **Data collection** | | | | | |
| Beamline | PETRA III P13 | ESRF ID30B | ESRF ID23-2 | ESRF ID30B | PETRA III P14 |
| Wavelength (Å) | 0.9763 | 0.9763 | 0.8731 | 0.9763 | 0.9763 |
| Space Group | I 222 | I 222 | I 222 | I 222 | I 222 |
| Unit cell<br>a b c (Å)<br>α β γ (°) | 63.15 98.46<br>140.62<br>90.0 90.0 90.0 | 62.98 92.12 139.85<br>90.0 90.0 90.0 | 63.56 99.08 140.620<br>90.0 90.0 90.0 | 62.69 98.90<br>139.59<br>90.0 90.0 90.0 | 63.12 98.21<br>140.10<br>90.0 90.0 90.0 |
| Resolution (Å) | 19.76 - 1.55 | 19.70 - 1.54 | 29.38 - 1.85 | 19.62 - 1.41 | 19.74 - 1.55 |

| | | | | | |
|---|---|---|---|---|---|
| | (1.59 - 1.55) | (1.58 - 1.54) | (1.90 - 1.85) | (1.45 - 1.41) | (1.59 - 1.55) |
| Unique reflections | 62272 | 64212 | 38279 | 83528 | 63211 |
| Multiplicity | 8.71 (8.22) | 6.27 (5.60) | 7.59 (7.71) | 6.26 (6.33) | 13.40 (10.88) |
| Completeness (%) | 97.5 (96.4) | 99.8 (100.0) | 99.9 (100.0) | 99.8 (99.8) | 99.3 (98.3) |
| $I$ / σ$I$ | 22.46 (2.50) | 15.11 (3.18) | 17.09 (3.51) | 12.85 (3.24) | 25.55 (3.13) |
| $R_{merge}$ | 0.062 (1.123) | 0.064 (0.502) | 0.083 (0.641) | 0.075 (0.605) | 0.055 (0.831) |
| $R_{meas}$ | 0.066 (1.199) | 0.070 (0.554) | 0.090 (0.688) | 0.082 (0.660) | 0.058 (0.872) |
| $CC_{1/2}$ | 100.0 (83.0) | 99.8 (90.1) | 99.9 (90.3) | 99.5 (85.5) | 100 (83.3) |
| **Refinement** | | | | | |
| $R_{work}$ / $R_{free}$ | 0.1708 / 0.1798 | 0.1618 / 0.1800 | 0.1588 / 1.893 | 0.1597 / 0.1754 | 0.1598 / 0.1742 |
| RMS bonds | 0.006 | 0.015 | 0.019 | 0.012 | 0.007 |
| RMS angles | 0.774 | 1.353 | 1.320 | 1.127 | 1.012 |
| Ramachandran | | | | | |
| favored (%) | 98.05 | 98.70 | 98.37 | 98.70 | 98.37 |
| allowed (%) | 1.95 | 1.30 | 1.63 | 1.30 | 1.63 |
| outliers (%) | 0.00 | 0.00 | 0.00 | 0.00 | 0.00 |
| Rotamer outliers (%) | 0.00 | 0.00 | 0.40 | 0.40 | 0.80 |
| Number of atoms | 2691 | 2674 | 2727 | 2745 | 2724 |
| Protein | 2364 | 2394 | 2363 | 2397 | 2374 |
| Ligands | 64 | 52 | 52 | 52 | 56 |
| Solvent | 263 | 228 | 312 | 296 | 294 |
| Average B-factor | 25.90 | 28.00 | 24.20 | 21.36 | 27.61 |
| Protein | 24.97 | 27.30 | 23.09 | 20.22 | 26.67 |
| Ligands | 20.80 | 23.31 | 19.87 | 15.52 | 22.07 |
| Solvent | 35.49 | 36.40 | 54.51 | 31.68 | 36.29 |

Values in parentheses correspond to the highest resolution shell.

### Supplementary table 2: List of strains used in this study.

| **Strain** | **Genotype** | **Source** |
|---|---|---|
| *E. coli* MG1655 | *E. coli* K-12 F−, λ−, *rph*-1 | Lab collection |
| *E. coli* SIJ488 | MG1655 Tn7::para-exo-beta-gam; prha-FLP; xylSpm-IsceI | Lab collection |
| DH5a | F− λ− Φ80*lac*ZΔM15 Δ(*lac*ZYA-*arg*F)U169 *deo*R *rec*A1 *end*A1 *hsd*R17(rK− mK+) *pho*A *sup*E44 *thi*-1 *gyr*A96 *rel*A1 | Lab collection |
| BL21 (DE3) | *F– ompT hsdSB (rB–, mB–) gal dcm (DE3)* | Lab collection |
| PZ | SIJ488 Δ*pfkA* Δ*pfkB* Δ*zwf* | |
| PZF | SIJ488 Δ*pfkA* Δ*pfkB* Δ*zwf* Δ*fsaA* Δ*fsaB* | |
| PZ mut 1 | SIJ488 Δ*pfkA* Δ*pfkB* Δ*zwf* plus mutations see Table 2 | This study |

| | | |
|---|---|---|
| PZ mut 2 | SIJ488 Δ*pfkA* Δ*pfkB* Δ*zwf* plus mutations see Table 2 | This study |
| PZ mut 3 | SIJ488 Δ*pfkA* Δ*pfkB* Δ*zwf* plus mutations see Table 2 | This study |
| PZ mut4 | SIJ488 Δ*pfkA* Δ*pfkB* Δ*zwf* plus mutations see Table 2 | This study |
| PZF mut 1 | SIJ488 Δ*pfkA* Δ*pfkB* Δ*zwf* Δ*fsaA* Δ*fsaB* plus mutations see Table 2 | This study |
| PZF mut 2 | SIJ488 Δ*pfkA* Δ*pfkB* Δ*zwf* Δ*fsaA* Δ*fsaB* plus mutations see Table 2 | This study |
| PZF mut 3 | SIJ488 Δ*pfkA* Δ*pfkB* Δ*zwf* Δ*fsaA* Δ*fsaB* plus mutations see Table 2 | This study |
| PZF mut 4 | SIJ488 Δ*pfkA* Δ*pfkB* Δ*zwf* Δ*fsaA* Δ*fsaB* plus mutations see Table 2 | This study |
| PZF mut 5 | SIJ488 Δ*pfkA* Δ*pfkB* Δ*zwf* Δ*fsaA* Δ*fsaB* plus mutations see Table 2 | This study |
| PZF mut 6 | SIJ488 Δ*pfkA* Δ*pfkB* Δ*zwf* Δ*fsaA* Δ*fsaB* plus mutations see Table 2 | This study |
| PZ mut 1 Δ*fruK* | SIJ488 Δ*pfkA* Δ*pfkB* Δ*zwf* Δ*fruK* plus mutations see Table 2 | This study |
| PZF mut 1 Δ*fruK* | SIJ488 Δ*pfkA* Δ*pfkB* Δ*zwf* Δ*fsaA* Δ*fsaB* Δ*fruK* plus mutations see Table 2 | This study |
| PZF+pORTMAGE-2 | SIJ488 Δ*pfkA* Δ*pfkB* Δ*zwf* | This study |
| PZF-K95H+pORTMAGE-2 | SIJ488 Δ*pfkA* Δ*pfkB* Δ*zwf* Δ*fsaA* Δ*fsaB* | This study |
| PZF-K95HN-Δ311:313+pORTMAGE-2 | SIJ488 Δ*pfkA* Δ*pfkB* Δ*zwf* Δ*fsaA* Δ*fsaB* | This study |

| | | |
|---|---|---|
| PZF-K95HN-Δ313:315+pORTMAGE-2 | SIJ488 Δ*pfkA* Δ*pfkB* Δ*zwf* Δ*fsaA* Δ*fsaB* | This study |
| PZF-K95HN-Δ306:308+pORTMAGE-2 | SIJ488 Δ*pfkA* Δ*pfkB* Δ*zwf* Δ*fsaA* Δ*fsaB* | This study |
| PZF-K95T+pORTMAGE-2 | SIJ488 Δ*pfkA* Δ*pfkB* Δ*zwf* Δ*fsaA* Δ*fsaB* | This study |
| PZF-K95T-G110S+pORTMAGE-2 | SIJ488 Δ*pfkA* Δ*pfkB* Δ*zwf* Δ*fsaA* Δ*fsaB* | This study |
| PZF-K95T-Q84R+pORTMAGE-2 | SIJ488 Δ*pfkA* Δ*pfkB* Δ*zwf* Δ*fsaA* Δ*fsaB* | This study |
| PZF-K95T-T34I+pORTMAGE-2 | SIJ488 Δ*pfkA* Δ*pfkB* Δ*zwf* Δ*fsaA* Δ*fsaB* | This study |
| PZF-K95S+pORTMAGE-2 | SIJ488 Δ*pfkA* Δ*pfkB* Δ*zwf* Δ*fsaA* Δ*fsaB* | This study |
| PZF-G110S+pORTMAGE-2 | SIJ488 Δ*pfkA* Δ*pfkB* Δ*zwf* Δ*fsaA* Δ*fsaB* | This study |
| PZF-G110A+pORTMAGE-2 | SIJ488 Δ*pfkA* Δ*pfkB* Δ*zwf* Δ*fsaA* Δ*fsaB* | This study |
| PZF-K95H | SIJ488 Δ*pfkA* Δ*pfkB* Δ*zwf* Δ*fsaA* Δ*fsaB* | This study |
| PZF-K95HN-Δ311:313 | SIJ488 Δ*pfkA* Δ*pfkB* Δ*zwf* Δ*fsaA* Δ*fsaB* | This study |
| PZF-K95HN-Δ313:315 | SIJ488 Δ*pfkA* Δ*pfkB* Δ*zwf* Δ*fsaA* Δ*fsaB* | This study |
| PZF-K95HN-Δ306:308 | SIJ488 Δ*pfkA* Δ*pfkB* Δ*zwf* Δ*fsaA* Δ*fsaB* | This study |
| PZF-K95T | SIJ488 Δ*pfkA* Δ*pfkB* Δ*zwf* Δ*fsaA* Δ*fsaB* | This study |
| PZF-K95T-G110S | SIJ488 Δ*pfkA* Δ*pfkB* Δ*zwf* Δ*fsaA* Δ*fsaB* | This study |
| PZF-K95T-Q84R | SIJ488 Δ*pfkA* Δ*pfkB* Δ*zwf* Δ*fsaA* Δ*fsaB* | This study |
| PZF-K95T-T34I | SIJ488 Δ*pfkA* Δ*pfkB* Δ*zwf* Δ*fsaA* Δ*fsaB* | This study |
| PZF-K95S | SIJ488 Δ*pfkA* Δ*pfkB* Δ*zwf* Δ*fsaA* Δ*fsaB* | This study |

| | | |
|---|---|---|
| PZF-G110S | SIJ488 Δ*pfkA* Δ*pfkB* Δ*zwf* Δ*fsaA* Δ*fsaB* | This study |
| PZF-G110A | SIJ488 Δ*pfkA* Δ*pfkB* Δ*zwf* Δ*fsaA* Δ*fsaB* | This study |
| PZFK* | SIJ488 Δ*pfkA* Δ*pfkB* Δ*zwf* Δ*fsaA* Δ*fsaB* Δ*fruK* plus mutations see Table 2 re-evolved for growth on glucose | This study |
| PZ +pCgFruK | SIJ488 Δ*pfkA* Δ*pfkB* Δ*zwf* Δ*fruK* | This study |
| PZ +pCgFruK-L29F | SIJ488 Δ*pfkA* Δ*pfkB* Δ*zwf* Δ*fruK* | This study |
| PZ +pCgFruK-G104A | SIJ488 Δ*pfkA* Δ*pfkB* Δ*zwf* Δ*fruK* | This study |
| PZF +pCgFruK | SIJ488 Δ*pfkA* Δ*pfkB* Δ*zwf* Δ*fsaA* Δ*fsaB* | This study |
| PZF +pCgFruK-L29F | SIJ488 Δ*pfkA* Δ*pfkB* Δ*zwf* Δ*fsaA* Δ*fsaB* | This study |
| PZF +pCgFruK-G104A | SIJ488 Δ*pfkA* Δ*pfkB* Δ*zwf* Δ*fsaA* Δ*fsaB* | This study |
| PZFK* +pCgFruK | SIJ488 Δ*pfkA* Δ*pfkB* Δ*zwf* Δ*fsaA* Δ*fsaB* Δ*fruK* plus mutations see Table 2 re-evolved for growth on glucose | This study |
| PZFK* +pCgFruK-L29F | SIJ488 Δ*pfkA* Δ*pfkB* Δ*zwf* Δ*fsaA* Δ*fsaB* Δ*fruK* plus mutations see Table 2 re-evolved for growth on glucose | This study |
| PZFK* +pCgFruK-G104A | SIJ488 Δ*pfkA* Δ*pfkB* Δ*zwf* Δ*fsaA* Δ*fsaB* Δ*fruK* plus mutations see Table 2 re-evolved for growth on glucose | This study |

| | | |
|---|---|---|
| *C. glutamicum* ATCC13032 | Wild-type (WT) | Lab Collection |
| PZ | ATCC13032 Δ*pfk* Δ*zwf* | This study |
| PZF | ATCC13032 Δ*pfk* Δ*zwf* Δ*fruk1* | This study |
| PZ mut 1 | ATCC13032 Δ*pfk* Δ*zwf* plus mutations see Table 3 evolved for growth on glucose | This study |
| PZ mut 2 | ATCC13032 Δ*pfk* Δ*zwf* plus mutations see Table 3 evolved for growth on glucose | This study |
| PZ mut 3 | ATCC13032 Δ*pfk* Δ*zwf* plus mutations see Table 3 evolved for growth on glucose | This study |
| PZ mut 4 | ATCC13032 Δ*pfk* Δ*zwf* plus mutations see Table 3 evolved for growth on glucose | This study |
